# An increase in surface hydrophobicity mediates chaperone activity in N-chlorinated proteins

**DOI:** 10.1101/2021.10.26.465855

**Authors:** Marharyta Varatnitskaya, Julia Fasel, Alexandra Müller, Natalie Lupilov, Yunlong Shi, Kristin Fuchs, Marco Krewing, Christoph Jung, Timo Jakob, Barbara Sitek, Julia E. Bandow, Kate S. Carroll, Eckhard Hoffmann, Lars I. Leichert

## Abstract

Under physiological conditions, *Escherichia coli* RidA is an enamine/imine deaminase, which promotes the release of ammonia from reactive enamine/imine intermediates. However, when modified by hypochlorous acid (HOCl), as produced by the host defense, RidA_HOCl_ turns into a potent chaperone-like holdase that can effectively protect the proteome of *E. coli* during oxidative stress. We previously reported that the activation of RidA’s chaperone-like function coincides with the addition of at least seven and up to ten chlorine atoms. These atoms are reversibly added to basic amino acids in RidA_HOCl_ and removal by reducing agents leads to inactivation. Nevertheless, it remains unclear, which residues in particular need to be chlorinated for activation. Here, we employ a combination of LC-MS/MS analysis, a chemo-proteomic approach, and a mutagenesis study to identify residues responsible for RidA’s chaperone-like function. Through LC-MS/MS of digested RidA_HOCl_, we obtained direct evidence of the chlorination of one arginine residue (and, coincidentally, two tyrosine residues), while other N- chlorinated residues could not be detected, presumably due to the instability of the modification and its potential interference with a proteolytic digest. Therefore, we established a chemoproteomic approach using 5-(dimethylamino) naphthalene-1-sulfinic acid (DANSO_2_H) as a probe to label N-chlorinated lysines. Using this probe, we were able to detect the N-chlorination of six additional lysine residues. Moreover, using a mutagenesis study to genetically probe the role of single arginine and lysine residues, we found that the removal of arginines R105 and R128 leads to a substantial reduction of RidAHOCl’s chaperone activity. These results, together with structural analysis, confirm that the chaperone activity of RidA is concomitant with the loss of positive charges on the protein surface, leading to an increased overall protein hydrophobicity. Molecular modelling of RidA_HOCl_ and the rational design of a RidA variant that shows chaperone activity even in the absence of HOCl further supports our hypothesis. Our data provide a molecular mechanism for HOCl-mediated chaperone activity found in RidA and a growing number of other HOCl-activated chaperones.

## INTRODUCTION

Phagocytosis is a crucial mechanism of our innate immune system used in the defense against and the elimination of pathogens, such as bacteria and fungi. Within the phagosome, a cellular compartment formed from the cytoplasmic membrane during phagocytosis, pathogens are exposed to a complex mixture of different reactive oxygen and nitrogen species, including superoxide radicals, hydrogen peroxide, peroxynitrite, and hypochlorous acid (for comprehensive reviews see Mortaz et al., 2018; Winterbourn et al., 2016).

Hypochlorous acid is produced by the heme enzyme myeloperoxidase from H_2_O_2_ and chlorine. HOCl is a highly reactive oxidizing and chlorinating agent and one of the most potent oxidants that exist in human cells (Albrich et al., 1981). It is known to cause a variety of modifications in virtually all cellular macromolecules including DNA (Prütz, 1996), lipids (Winterbourn et al., 1992; Carr et al., 1996; Deborde and von Gunten, 2008), carbohydrates (McGowan and Thompson, 1989) and proteins. Amino acid modifications caused by HOCl include oxidation of cysteine and methionine residues and chlorination of tyrosine, lysine, arginine, and histidine (Hawkins et al., 2003; Pattison et al., 2012). Often these modifications are irreversible, leading to protein misfolding and aggregation (Hawkins and Davies, 1999; Winter et al., 2008; Müller et al., 2014), loss of function, and eventual degradation by proteolytic enzymes (Hawkins and Davies, 1999). Thus, modification of proteins by oxidative stressors and HOCl, in particular, is mostly detrimental to their function. However, in some cases, oxidative modifications can also regulate and activate functions of some proteins to help the cell overcome those oxidative stress conditions. Long-established among the oxidative modifications that can lead to protein activation are reversible oxidations of the side chain of cysteines. Prime examples are transcription factors such as OxyR (Storz et al., 1990; Zheng et al., 1998) or the redox-regulated chaperone Hsp33 (Jakob et al., 1999) in *E. coli*. Both proteins become activated upon the formation of disulfide bonds and, conversely, reduction of these disulfides leads to their inactivation. More recently we discovered that N-chlorination of basic amino acid side chains, such as lysine and arginine, is another *in vivo*-reversible modification that mediates the activation of a chaperone-like holdase function in the bacterial protein RidA. After treatment with HOCl or monochloramine, we observed an approximately 60 % reduction in the amino group content accompanied by an increase in RidA’s hydrophobicity. Consistent with this finding, mass spectrometry (MS) analysis revealed the presence of multiple chlorinated species (Müller et al., 2014). This activation of RidA is independent of the oxidation status of its sole cysteine and is fully reversible by DTT, ascorbic acid, glutathione, and the thioredoxin system.

Other proteins, like the bacterial protein CnoX have been found to use a similar mechanism. After activation with HOCl, CnoX, similar to RidA, turns into an effective chaperone-like holdase that can bind a variety of substrate proteins and prevent their aggregation (Goemans et al., 2018). Moreover, CnoX has a thioredoxin domain and can protect its substrates from irreversible thiol oxidation by forming mixed disulfides with them (Goemans et al., 2018). But not only bacterial proteins are affected by N- chlorination: recently we discovered that human plasma proteins, such as human serum albumin, are activated by N-chlorination and act not only as chaperones to protect damaged host proteins but can also activate immune cells promoting their production of ROS (Ulfig et al., 2019) and mediating their antigen processing (Ulfig et al., 2021).

In this study, we identified the residues facilitating RidA’s HOCl-driven activation mechanism. Our data suggest that chlorination of basic amino acids by HOCl is not a process that targets one or two crucial amino acids that act as the “switch” affecting the whole protein. Instead, the N-chlorination of multiple residues changes the electrostatic potential of the molecular surface of RidA allowing it to interact with unfolded substrate proteins.

## RESULTS

### Direct identification of chlorinated sites in RidA after treatment with HOCl revealed chlorination of one arginine and two tyrosines

We previously reported that the enamine/imine deaminase RidA from *E. coli* turns into a potent protein holdase upon N-chlorination of its lysine and arginine residues. RidA, modified in this way (RidA_HOCl_), can protect other cellular proteins from unfolding due to chlorine stress. Previous mass spectrometric analysis of undigested HOCl-treated RidA showed that the reversible addition of at least 7 and up to 10 chlorine atoms to amino acid residues is concomitant with chaperone-like holdase activity in this protein (Müller et al., 2014). However, it remains unknown, which residues are modified and which of those are required for activation. If we consider arginine, histidine, and lysine as potential targets for a reversible N-chlorination, RidA has 15 possible N-chlorination sites: 5 arginines, 1 histidine, 8 lysines, and the protein’s N-terminus (Figure 1).

**Figure 1.**
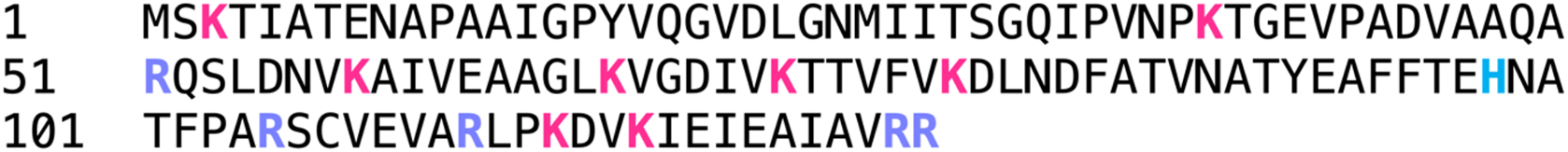
The amino acid sequence of the *E. coli* protein RidA (Uniprot entry number: P0AF93). Lysines (K) are highlighted in pink, arginines (R) in purple, and histidine (H) in blue.

To elucidate which amino acids are affected by chlorination, we first performed a mass spectrometric analysis of N-chlorinated RidA after proteolytic digest, specifically searching for peptides having an added mass due to chlorination. Among the above-mentioned 15 possible N-chlorination sites, we were only able to identify R51 to be chlorinated in any of the three replicates after digest with trypsin (Table 1, Figure 2). The monoisotopic mass of the chlorinated peptide containing R51 was ∼ 33.96 Da heavier than the unmodified peptide, corresponding to the monoisotopic mass of a chlorine atom minus the monoisotopic mass of the hydrogen atom it replaced. The same tell-tale mass difference could be observed in 9 fragment y-ions containing R51 (Figure 2C). Additionally, we found two chlorinated peptides containing the two tyrosines present in RidA (Y17, Y91) (Table 1). Tyrosine chlorination by HOCl is an irreversible modification that occurs with a much slower rate than the chlorination of lysine, histidine and arginine (Hawkins et al., 2003). But once formed, chlorotyrosine is highly stable (Winterbourn and Kettle, 2000; Hendrikje Buss et al., 2003). Our previous experiments showed that the 7 to 10 added chlorine atoms observed in full-length MS are virtually all removable by DTT, inconsistent with chlorinated tyrosine residues. We thus concluded that our finding is, due to the high stability of the modification in question, probably of auxiliary nature with no functional relevance.

**Figure 2.**
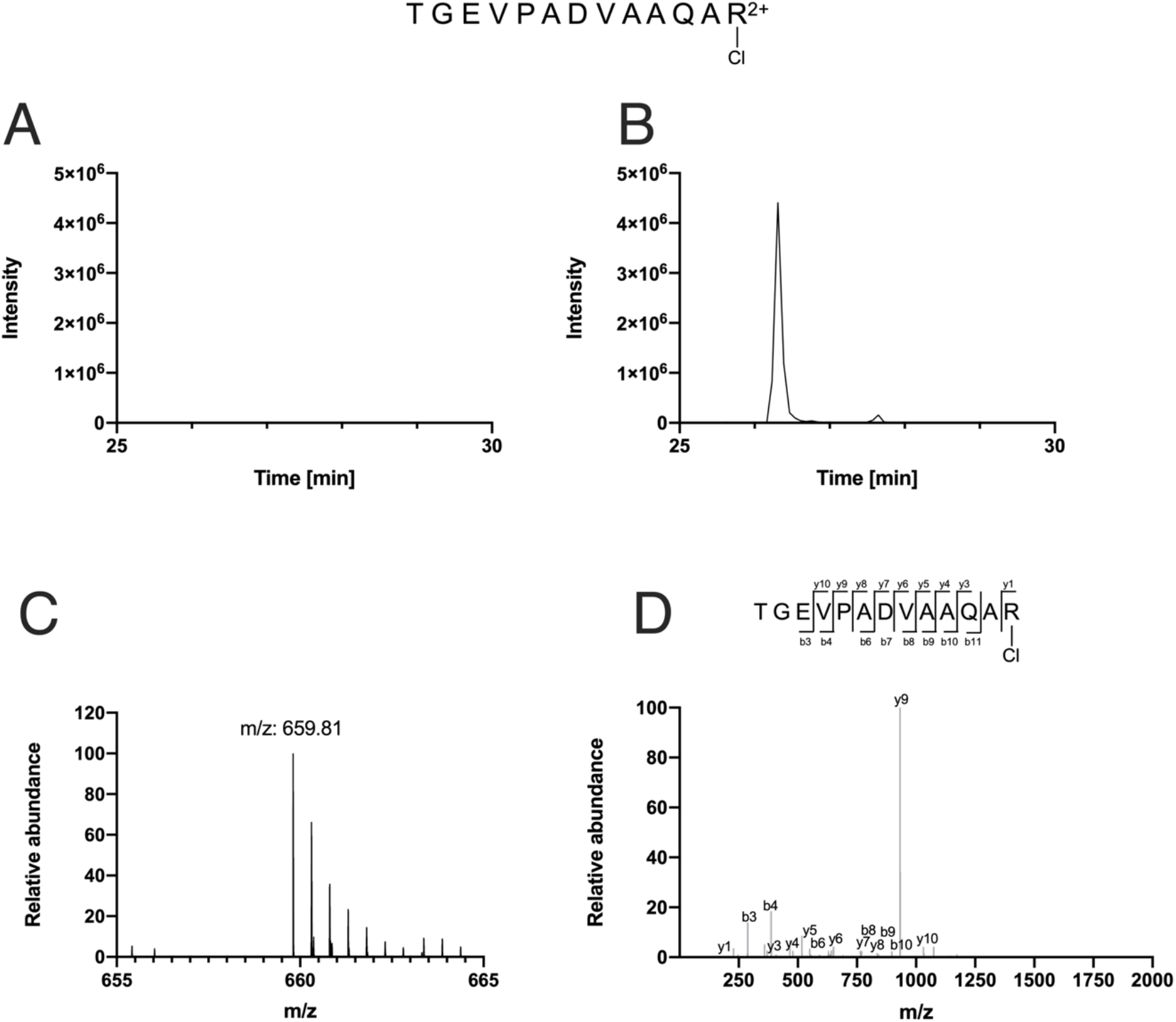
Arginine R51 is chlorinated upon treatment with HOCl. **(A, B)** Extracted ion chromatograms (XIC) of m/z 659.81, corresponding to the mass of the N-chlorinated peptide TGEVPADVAAQAR51 at a retention time from 25 to 30 minutes. A tryptic digest of untreated RidA produces no ion with the respective m/z **(A)**, while RidA treated with a 10-fold molar excess of HOCl shows a peak at 26.265 min **(B)**. **(C)** Primary MS spectrum of the N- chlorinated arginine-containing peptide TGEVPADVAAQAR51. **(D)** Corresponding fragment spectrum of the N- chlorinated peptide contains signals corresponding to 9 y-ions containing the chlorine atom.

**Table 1.**
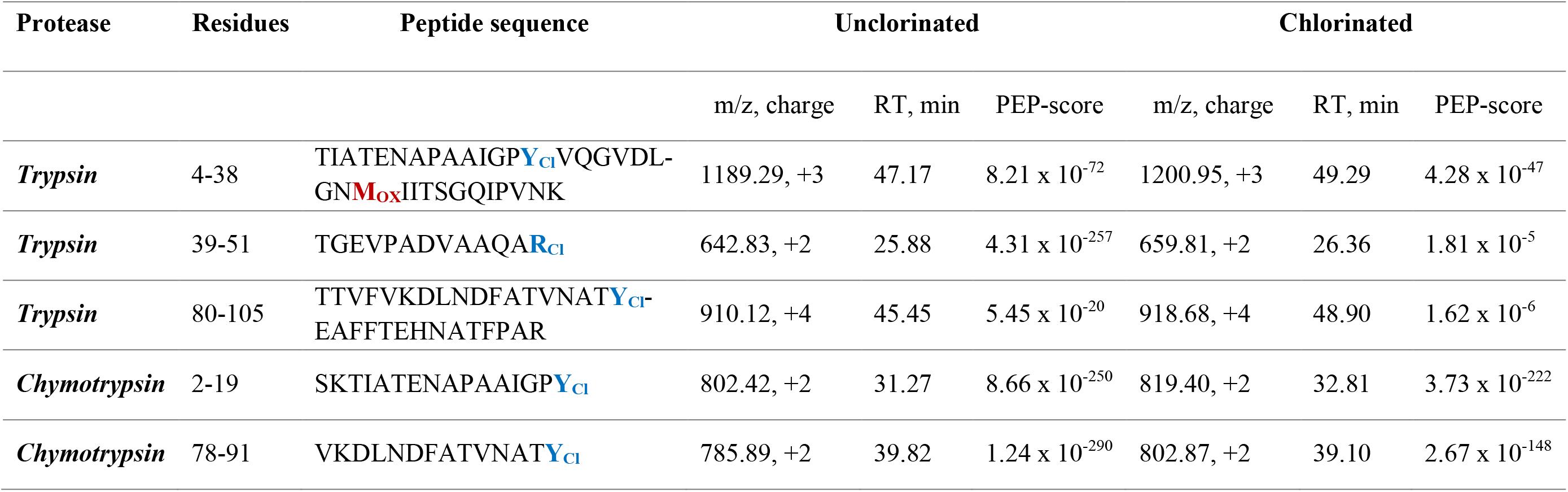
Retention times (RT) and properties of unchlorinated and chlorinated peptides identified by LC-MS/MS after tryptic or chymotryptic digest of HOCl-treated RidA. Methionine sulfoxidation is highlighted in red, and possible sites of chlorination are labeled blue. The PEP-score (posterior error probability) calculated by MaxQuant indicates the probability that a peptide was falsely identified (Käll et al., 2008; Cox and Mann,

We suspected that the low number of identified N-chlorinated residues in our experiment could be caused by our use of trypsin. Trypsin, the protease most commonly used for the MS analysis of proteins, cleaves proteins after lysine and arginine, the very residues which we suspected to be modified by HOCl. As chlorination of these residues might interfere with trypsin’s ability to recognize them, we additionally prepared a digest with chymotrypsin. Nevertheless, we were still not able to detect more N-chlorinated residues in our chymotryptic digest in any replicate but only re-identified chlorinated tyrosines Y17 and Y91 (Table 1).

### DANSO_2_H, a novel proteomic probe for N-chlorinated lysine residues in RidA_HOCl_

Our previous results showed that at least seven and up to ten residues become N-chlorinated in RidA, when it is active as a chaperone. Thus, we were dissatisfied with our limited ability to detect N- chlorinated amino acids using mass spectrometry of proteolytic digests of HOCl-treated RidA. This inability might be due to the inherent instability and high reactivity of N-chloramines. Therefore, we devised a way to stably modify N-chlorinated amino acids.

To this end we utilized dansyl sulfinic acid (5-(dimethylamino)naphthalene-l-sulfinic acid, DANSO_2_H), a derivative of the well-characterized reagent dansyl chloride (5-dimethy1amino)naphthalene-1-sulfonyl chloride, DANSCl), which has been used for a long time to derivatize amines, amino acids, and proteins. It reacts with free amino groups and forms stable, highly fluorescent sulfonamides, that can be separated by HPLC and detected by mass spectrometry. In proteins, DANSCl usually reacts only with the free amino group of lysine and the N-terminus (Hsieh and Matthews, 1985) (Figure 3A). Unlike DANSCl, DANSO_2_H is not reactive towards unmodified amino groups. Instead, it reacts with monochloramines, forming the same sulfonamide product as the reaction of DANSCl with corresponding unmodified amines (Figure 3C). This provides us with a method for the selective labeling of N-chlorinated lysine residues. DANSO_2_H was synthesized from DANSCl by a reaction with sodium sulfite (Scully et al., 1984). The synthesized molecule had a purity of 95 %. The only contamination present was dansyl sulfonic acid (DANSO_3_H) (Figure S2), which is chemically inert and does react neither with amino groups nor N-chlorinated residues (Scully et al., 1984). The synthesized DANSO_2_H was then used to modify RidA_HOCl_. DANSCl was used as a positive control to modify untreated RidA (RidA_UT_), and, as a negative control, DANSO_2_H was incubated with RidA_UT_. As expected, after treatment with DANSCl, RidA_UT_ showed the typical fluorescence of dansyl-modified proteins (Figure 3B). The same fluorescence albeit to a lesser extent was observed in DANSO_2_H-treated RidA_HOCl_ (Figure 3D). The ∼30 % lower fluorescence intensity is consistent with our previous finding that only up to 10 out of 15 potential chlorination sites are chlorinated in fully active RidA_HOCl_. RidA_UT_ treated with DANSO_2_H showed virtually no fluorescence, demonstrating the specificity of our probe (Figure 3D).

**Figure 3.**
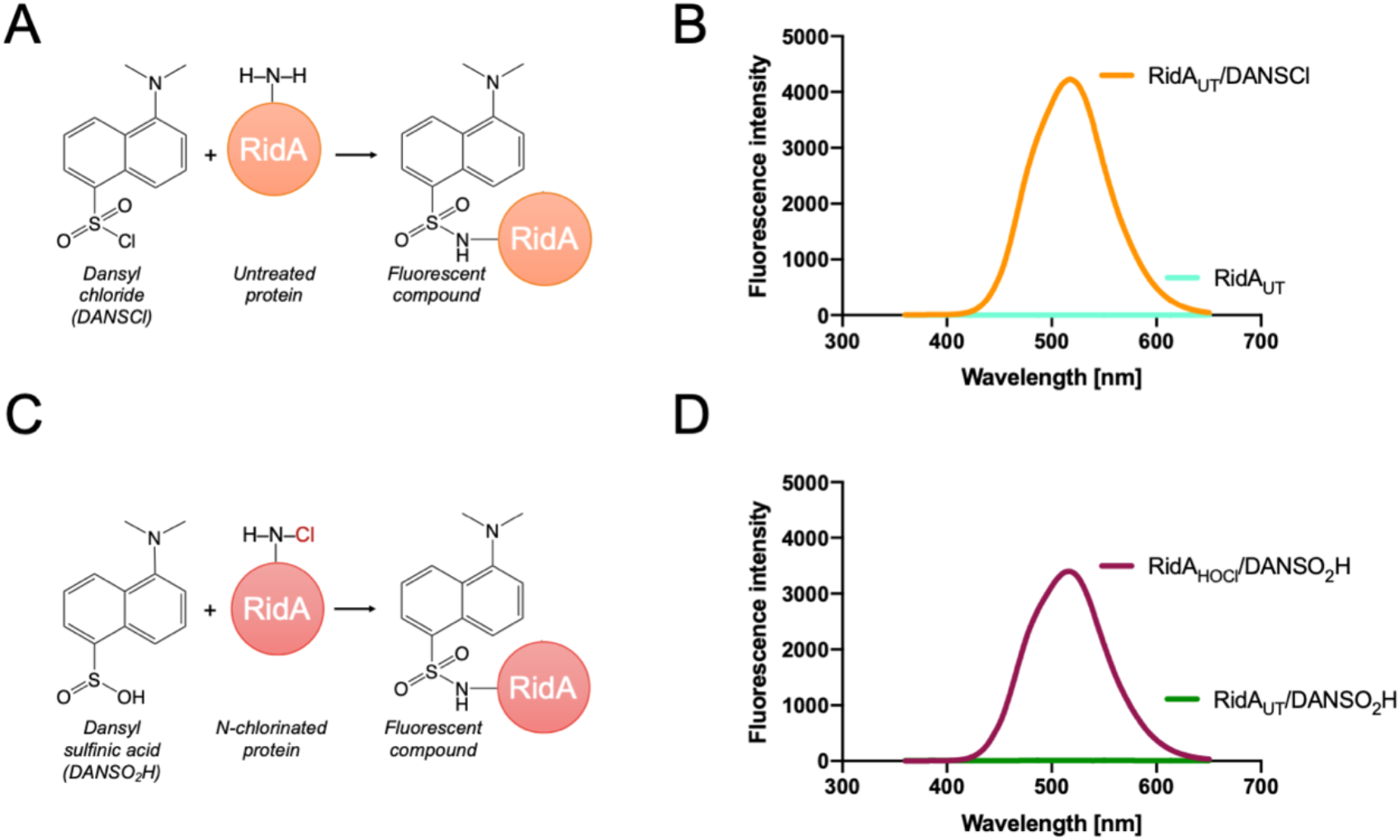
Modification of RidA with dansyl-derived compounds. **(A)** Dansyl chloride (DANSCl) reacts with an exemplary amino group in RidA forming a “dansylated” fluorescent sulfonamide. **(B)** Dansylation of proteins can be detected using fluorescence spectroscopy at Ex/Em: 340/500 nm. The reaction of RidAUT with a 10-fold molar excess of DANSCl yields a fluorescent protein (orange spectrum), while the protein by itself shows no fluorescence (teal spectrum). **(C)** Dansyl sulfinic acid (DANSO2H) is a derivative of DANSCl and reacts specifically with N-chlorinated amino groups forming the same “dansylated” fluorescent product found in panel (A). **(D)** HOCl-treated RidA (RidAHOCl) reacts with a 10-fold molar excess of DANSO2H producing a fluorescent peak at the expected excitation/emission wavelengths (purple spectrum). Virtually no fluorescent signal was observed with DANSO2H-treated RidAUT (green spectrum).

**Figure 4.**
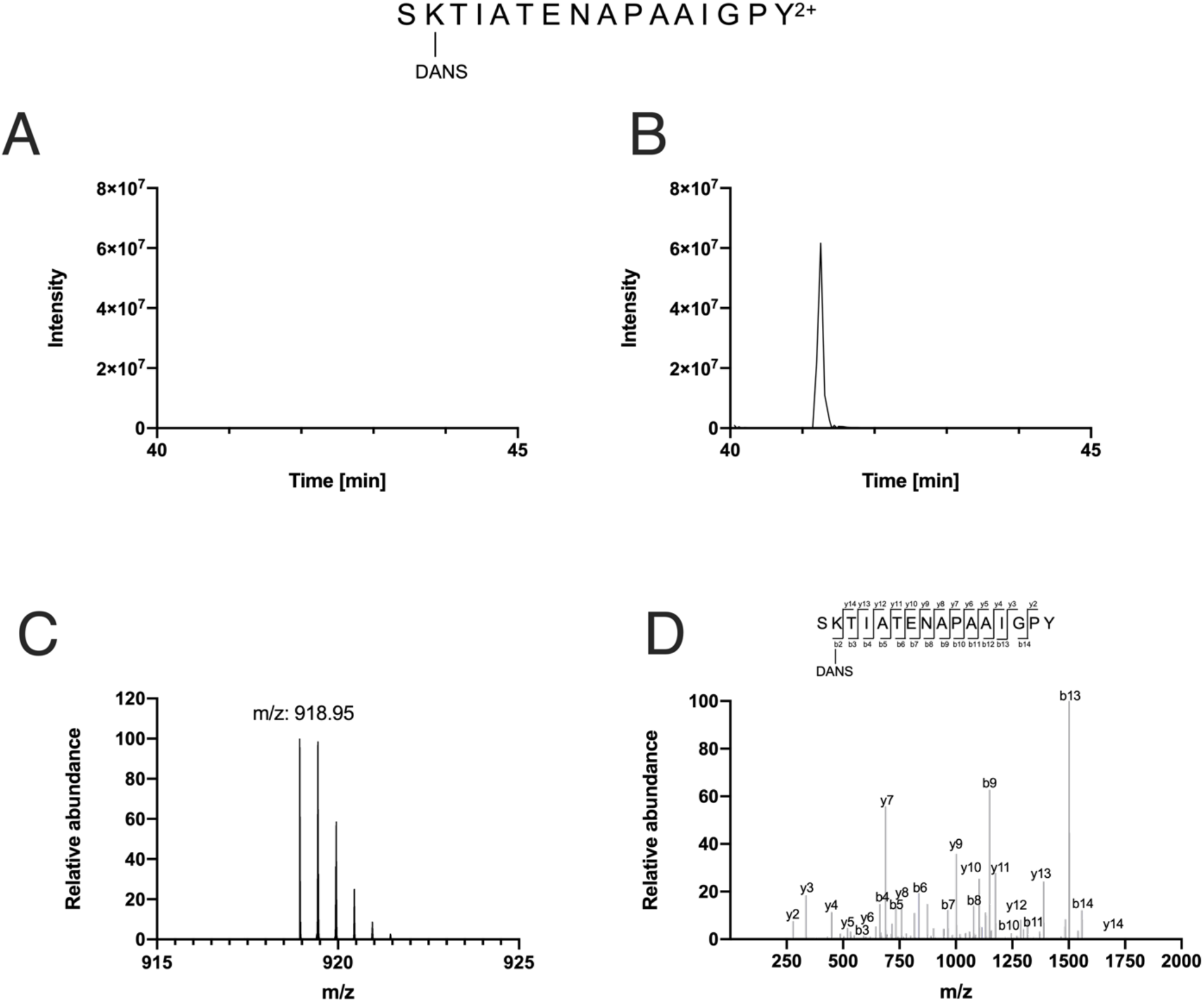
LC-MS/MS analysis of RidAHOCl after treatment with DANSO2H and subsequent chymotryptic digest. **(A, B)** Extracted ion chromatogram (XIC) of m/z 918.95, corresponding to the mass of dansylated peptide SK3TIATENAPAAIGPY at retention times from 40-45 min. **(A)** In the sample derived from DANSO2H-treated RidAUT, no ion at this m/z was observed. **(B)** A peak at 41.373 min corresponding to the mass of the dansylated peptide could be observed in DANSO2H-treated RidAHOCl. **(C)** Primary MS spectrum of the DANSO2H-modified peptide SK3TIATENAPAAIGPY found in the digest derived from DANSO2H-treated RidAHOCl, at an m/z of 918.95. **(D)** MS/MS analysis revealed that the modified amino acid is lysine K3 at position 2 of the chymotryptic peptide with one y-ion and thirteen b-ions showing the mass shift associated with dansylation.

### N-chlorinated lysines can be identified after labeling of RidA_HOCl_ with DANSO_2_H

Since the fluorescence spectroscopic experiments showed promising results, we then digested the DANSO_2_H-treated RidA_HOCl_ with chymotrypsin and performed LC-MS/MS analysis. In total, six lysines (K3, K38, K67, K79, K115, K118) were consistently identified as dansylated in all tested samples of DANSO_2_H-treated RidA_HOCl_ (Table 2). Masses corresponding to these dansylated peptides were not observed in RidA_UT_. Unfortunately, arginines seem not to be modified by DANSO_2_H, consistent with DANSCl’s preference for free amino groups. Dansylated peptides have a mass difference to the unmodified peptide of ∼ 233.05 Da, which equals the mass of DANSO_2_H minus the monoisotopic mass of two hydrogen atoms. The dansyl-modification changes the chemical property of a peptide making it more hydrophobic, causing it to elute at significantly higher acetonitrile (ACN) concentrations in the LC. For instance, unmodified peptide harboring K3 eluted at approx. 18 % ACN, whereas the corresponding dansyl-modified peptide eluted at approx. 27 % ACN (Table 2)

**Table 2.**
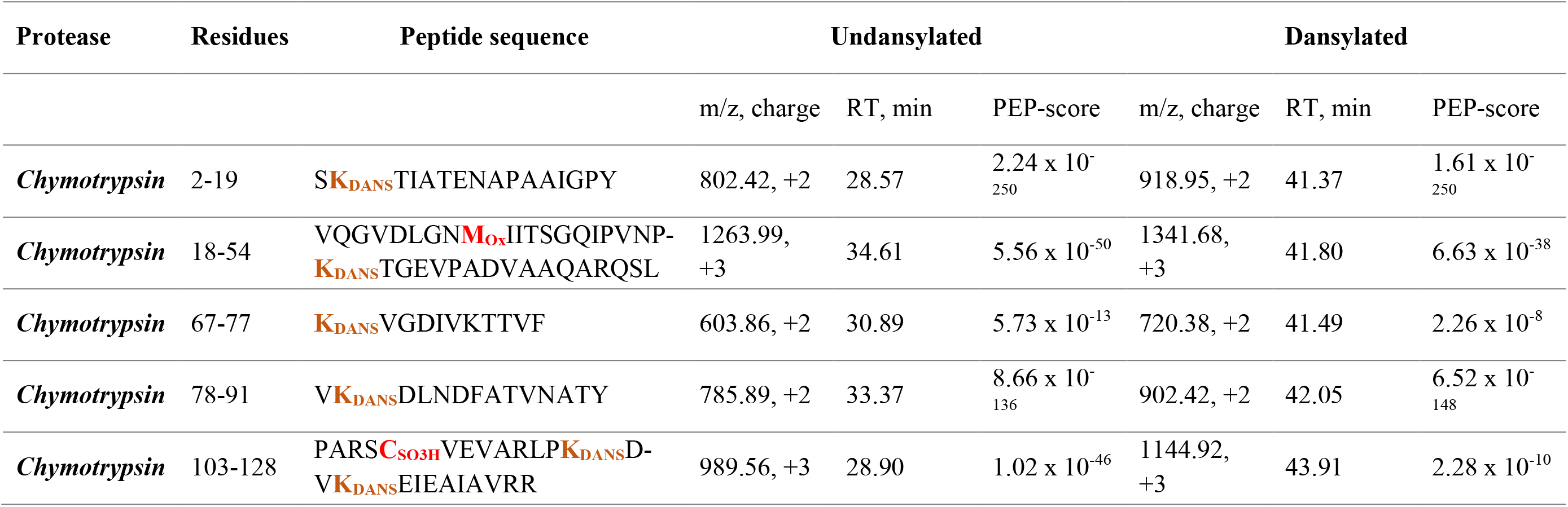
Retention times (RT) and properties of peptides derived from dansyl sulfinic acid. (DANSO2H)-labeling of HOCl-treated RidA after chymotryptic digest and LC-MS/MS. Methionine sulfoxidation and cysteine sulfonic acid modifications are highlighted in red, and lysines that are modified with dansyl sulfonic acid are labeled orange. The PEP-score calculated by MaxQuant indicates the probability that a peptide was falsely identified (Käll et al., 2008; Cox and Mann, 2008)

No dansylated amino acids were found in a RidA_UT_ sample treated with DANSO_2_H. With this novel chemoproteomic method we were able to gather direct evidence for the N-chlorination of six lysines in the active RidA_HOCl_ chaperone-like holdase.

### Mutagenesis reveals two arginines that play an important role in the activation of RidA’s chaperone activity

Since DANSO_2_H is most likely not suitable for the detection of N-chlorinated arginine residues, we used a mutagenesis approach to test if individual amino acids have a particular impact on the chaperone function of N-chlorinated RidA. We thus engineered a total of 13 RidA variants with every single lysine or arginine residue exchanged to a serine.

These 13 RidA variants were expressed and purified from *E. coli* BL21(DE3) and their chaperone activity after treatment with HOCl was then determined. Activity was tested in a 4-fold excess over citrate synthase. The activity of the variants was compared against wild-type RidA_HOCl_ under the same conditions. The activity of most of HOCl-treated mutants was not affected in a significant way by the exchange of a single N-containing amino acid to serine, when compared to HOCl-treated wild-type protein (Figure 5). However, two variants lacking specific arginines (R105S and R128S) showed a significantly reduced chaperone activity when compared to HOCl-treated wild-type RidA (RidA_WT_).

**Figure 5.**
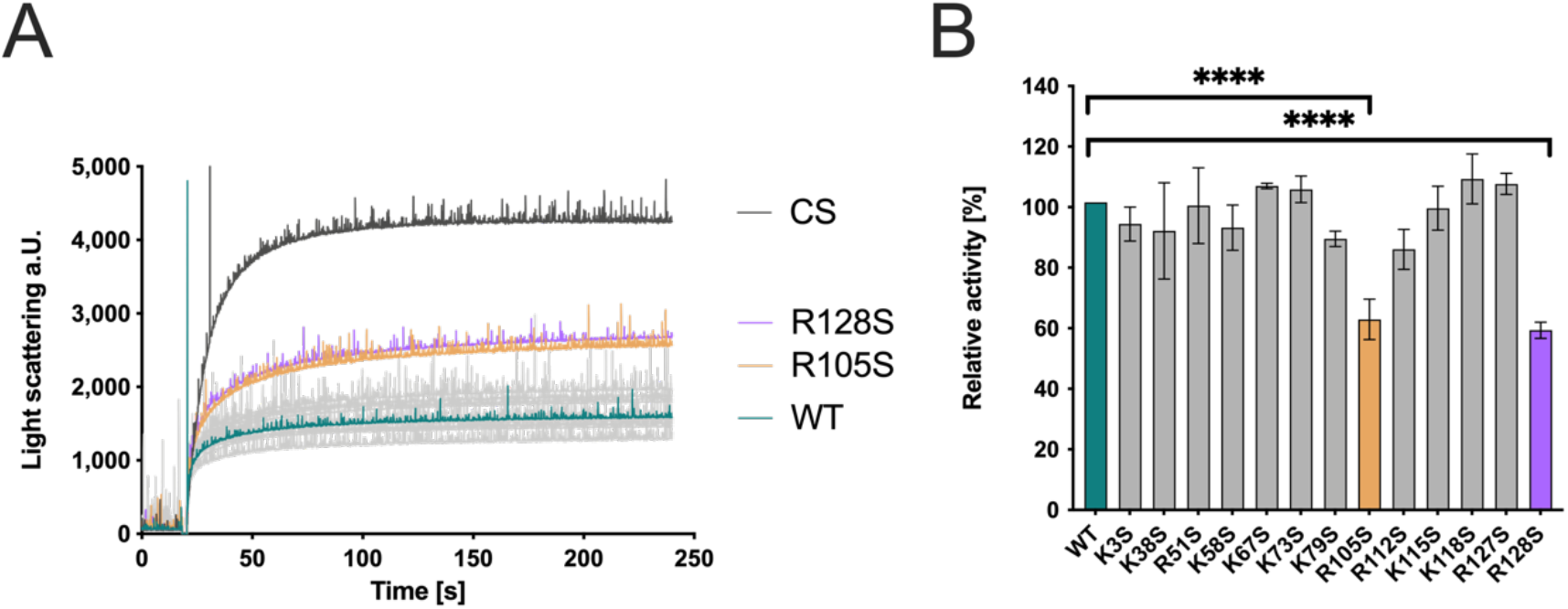
Chaperone activity of RidA variants lacking individual lysine or arginine residues. Chaperone activity was tested in an aggregation assay with citrate synthase. **(A)** HOCl-treated RidAWT (teal) strongly inhibits aggregation of chemically denatured citrate synthase when compared to untreated control (black), as measured by light scattering at 360 nm. All variants, except for R105S (yellow) and R128S (violet), showed activity similar to RidAWT. **(B)** Bar graph data represents means and standard deviations from three independent experiments. Differences in chaperone activity of RidAWT between the variants were analyzed using a one-way ANOVA with Tukey’s comparison test (****, p < 0.0001). The activity of RidAWT was set to 100 %, and all the data are presented in correlation to this control.

### The concomitant exchange of R105 and R128 does not further decrease chaperone activity

Since an exchange of arginine 105 or 128 resulted in decreased chaperone activity, a variant harboring both mutations (R105S_R128S) was constructed to investigate if a synergistic effect can be observed. The variants were then also treated with HOCl and used in an 8-fold excess over citrate synthase. However, the chaperone activity of the variant R105S_R128S remained at approximately the same level as the single exchange variants and a further decrease of chaperone activity was not observed (Figure 6).

**Figure 6.**
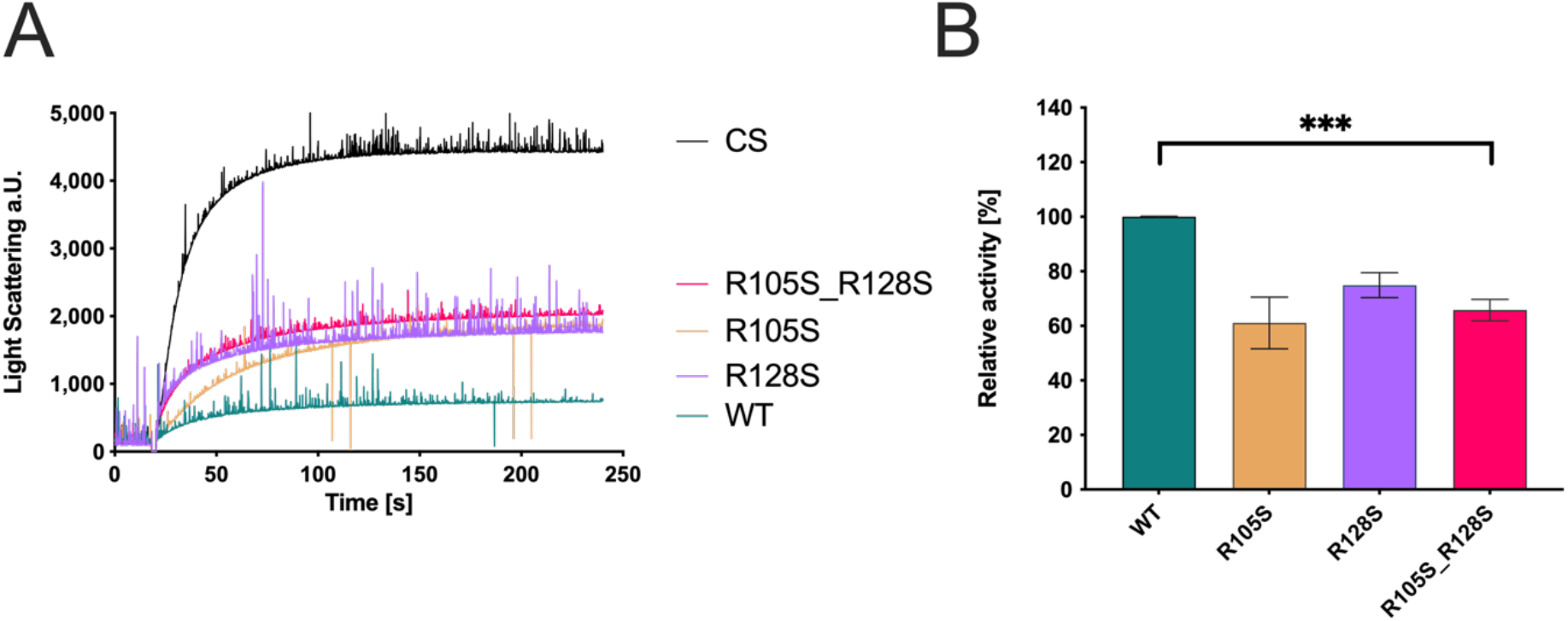
Chaperone activity of RidA double exchange variant R105S_R128S. Chaperone activity was tested in an aggregation assay with citrate synthase. **(A)** HOCl-treated RidAWT (teal) strongly inhibits aggregation of chemically denatured citrate synthase compared to untreated control (black), as measured by light scattering at 360 nm. The concomitant exchange of R105 and R128 against serine (pink) does not further decrease the chaperone activity of RidA and is comparable to single amino acid exchange variants (R105S (yellow) and R128S (violet)). **(B)** Bar graph data represents means and standard deviations from three independent experiments. Differences in chaperone activity between the variants were analyzed using a one-way ANOVA with Tukey’s comparison test (***, p < 0.001). The activity of RidAWT was set to 100 %, and all the data are presented in correlation to this control.

### The activation of RidA’s chaperone-like holdase function depends on an overall change of the molecules electrostatic surface

Summed up, our proteomic, chemo-proteomic and mutagenesis studies suggest that no single amino acid acts as a discrete “switch”, but rather the modification of multiple N-containing amino acids leads to the activation of RidA’s chaperone-like holdase function. Even arginines 105 and 128, whose individual exchanges to the inert amino acid serine had the largest effect on activatability, did not show synergistic effects when both were removed. Chaperones are known for surface patches that enable them to interact with hydrophobic regions of unfolded proteins. Indeed, our previous experiments showed that RidA_HOCl_ did bind a hydrophobic dye much better than RidA_UT_ (Müller et al., 2014). In order to understand how N-chlorination of basic amino acids changes the surface properties of RidA, we predicted the electrostatic surface potential of RidA, using the known X-Ray structure of RidA (Volz, 2008) (Figure 7A). We then computationally substituted lysine and arginine residues with their chlorinated counterparts within the structure data set. Using a customized force field, we were then able to predict the electrostatic surface potential of an N-chlorinated RidA molecule as well (Figure 7B). Strikingly, the electrostatic surface potential shifted towards a more negative potential, more or less over the complete surface of the molecule. We thus concluded that the loss of positive electrostatic surface potential is the underlying molecular reason for RidA_HOCl_’s chaperone-like properties.

**Figure 7.**
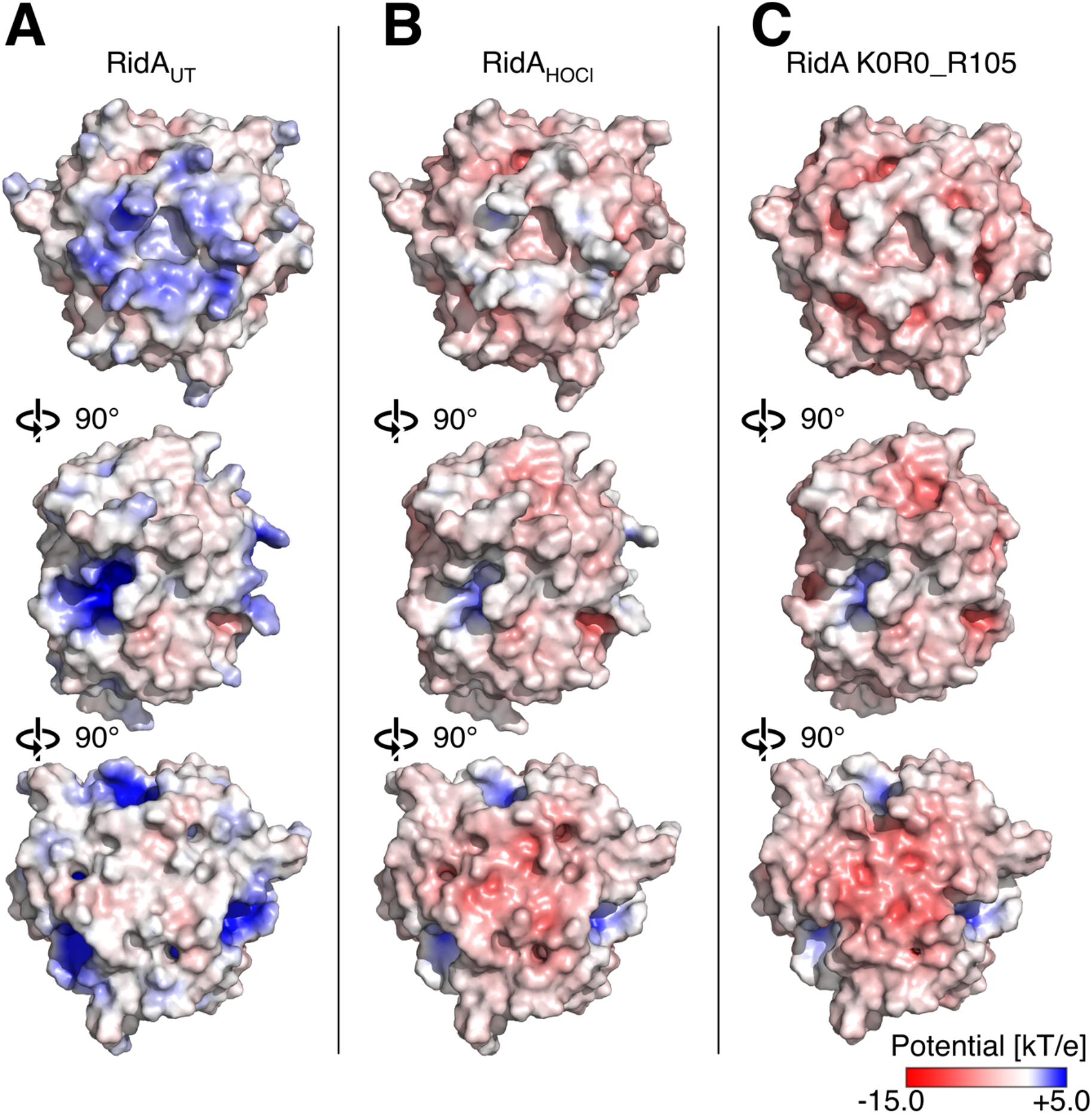
Calculated electrostatic surface potentials of *E. coli* RidAUT (Uniprot entry: P0AF93, PDB entry: 1QU9), RidAHOCl, and the K0R0_R105 variant. The C-terminal R128 was added to the RidA structure, as it was only partially resolved in the crystal structure. **(A)** Electrostatic surface visualization of wild-type RidA. The calculation was performed using the Advanced Poisson-Boltzmann Solver in PyMol using the AMBER force field, as described in the materials and methods section. The coloring scale was chosen from -15 to +5 kT/e according to the legend. Middle panel: the molecule rotated 90 degrees around the x-axis in comparison to the upper panel. Lower panel: the molecule was rotated 90 more degrees around the x-axis. **(B)** Electrostatic surface visualization of RidAHOCl using PyMOL. The charge-bearing proton was removed from all lysine and arginine residues, except for R105, and one of the hydrogens was then replaced by a chlorine atom using bond geometries derived from model compounds. The force field (partial charges of respective N and Cl atoms as well as the van der Waals radius of Cl) was adjusted accordingly. **(C)** Electrostatic surface visualization of K0R0_R105. Amino acid exchanges were introduced in RidA’s structure using PyMol’s “Mutagenesis” function.

### An engineered variant of RidA mimicking RidA_HOCl_’s surface potential is an active chaperone without HOCl treatment

To test our hypothesis, we decided to engineer a variant of RidA that mimics RidA_HOCl_’s surface potential. To this end, we wanted to mutate all lysine and arginine residues in RidA to more “electroneutral” amino acids. In order to select amino acids that would not lead to a disruption of RidA’s structure, we performed a multiple sequence alignment to direct our choice of suitable amino acids. For our mutagenesis, we selected amino acids that occur at the respective position and are mostly uncharged at physiological pH. With the exception of the invariant R105, the arginine in the active deaminase site of RidA, we were able to find a suitable amino acid at all positions. This resulted in a variant, which we termed R0K0_R105 (Figure 8).

**Figure 8.**
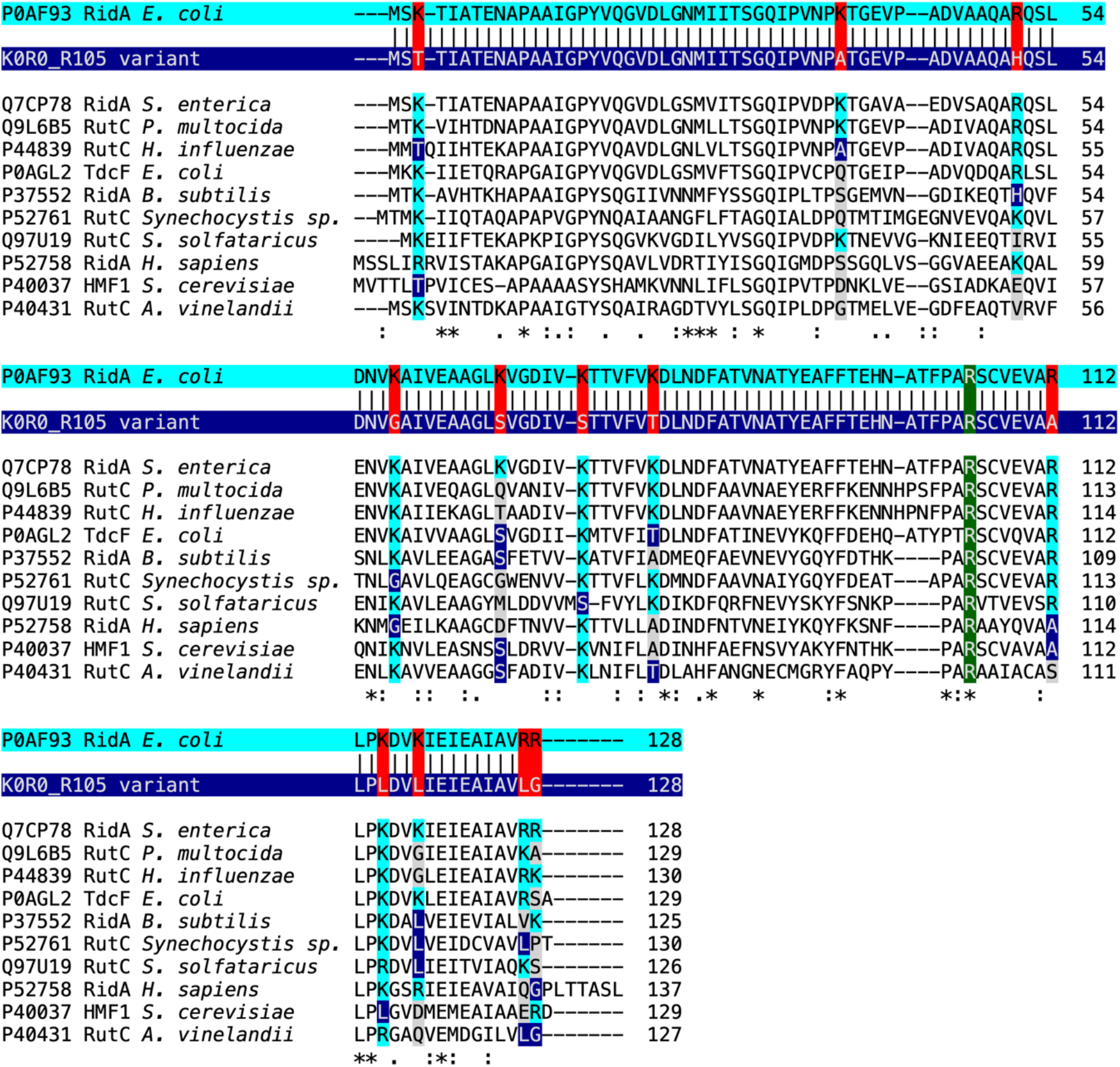
Sequence alignment of wild-type RidA (P0AF93 RidA *E. coli*) with the R0K0_R105 variant and RidA homologs. In the K0R0_R105 variant all lysine and arginine residues were replaced with more electroneutral residues based on amino acids found at those positions in RidA homologs, with the exception of the active-site arginine R105, where no replacement amino acid was found. The order of sequences is based on overall amino acid identity, ranging from 93.75 % *(S. enterica*) to 42.19 % (*A. vinelandii*). Arginine and lysine residues found at the position of the replaced amino acids are highlighted in cyan in the alignment, while amino acids chosen for replacement are highlighted in blue. Other amino acids at those positions are highlighted in grey. The invariant arginine R105 is highlighted in green.

The predicted electrostatic surface potential of this R0K0_R105 variant showed a high similarity to the predicted surface potential of N-chlorinated RidA_HOCl_ (Figure 7C).

The K0R0_R105 variant was then expressed and purified from *E. coli* BL21(DE3) and its chaperone activity in comparison to RidA_UT_ and RidA_HOCl_ was determined. As predicted, the RidA variant K0R0_R105 showed already potent chaperone-like activity without HOCl pre-treatment, and treatment with HOCl did not increase its chaperone activity significantly (Figure 9). Overall, the activity of K0R0_R105 was indeed comparable to HOCl-treated RidA, supporting our hypothesis.

**Figure 9.**
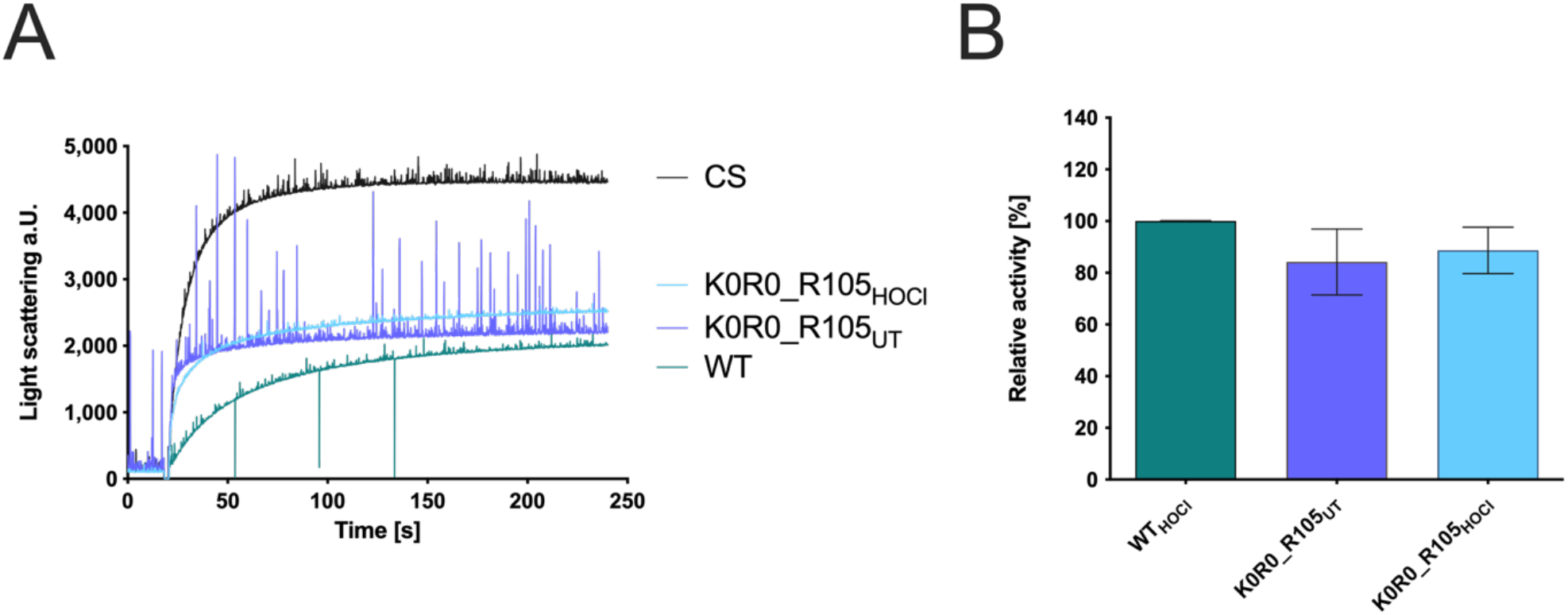
Chaperone activity of the RidA variant K0R0_R105. Chaperone activity was tested in an aggregation assay with citrate synthase. **(A)** HOCl-treated RidAWT (teal) strongly inhibits aggregation of chemically denatured citrate synthase compared to untreated control (black), as measured by light scattering at 360 nm. The variant K0R0_R105 (purple) is fully active as a chaperone without any HOCl treatment, and N-chlorination of this variant (light blue) does not influence its chaperone activity. **(B)** Bar graph data represents means and standard deviations of three independent experiments. Difference in chaperone activity between the untreated and HOCl-treated K0R0_R105 was analyzed using a one-way ANOVA with Tukey’s comparison test. The activity of RidAWT was set to 100 %, and all data are presented in correlation to this control.

Taken together, our results suggest that chlorination of basic amino acids by HOCl is not a process that targets one or two crucial amino acids that act as a “switch” that affects the whole protein. Instead, the N-chlorination of multiple residues generates a more negatively charged and hydrophobic molecular surface on RidA, allowing for the interaction with unfolded client proteins. This loss of positive surface charge through N-chlorination is a plausible mechanism for the chaperone-like switch that occurs in a growing number of proteins in response to exposure to HOCl and other reactive chlorine species.

## DISCUSSION

After reacting with HOCl or monochloramine, RidA turns into a potent chaperone-like holdase that can bind client proteins. The underlying mechanism was proposed to be the N-chlorination of lysine and arginine residues (Müller et al., 2014). Since then, several other proteins have been discovered that seem to switch to a chaperone-like function upon N-chlorination of basic amino acids. However, it was still unclear if the chlorination of one or several specific residues is decisive for chaperone activity of RidA or rather a general modification of multiple amino acid side chains that affects the overall surface of the protein.

Four proteinogenic amino acids are known to be prone to chlorination (Peskin and Winterbourn, 2001). First, lysines have an amino group on their side chain that can react with HOCl, forming monochloramines. The guanidino group of arginine can also be N-chlorinated. These modifications are thought to be reversed by ascorbate. The side chain of histidine can also react with HOCl, forming a short-lived chloramine. Nevertheless, this chloramine has been shown to be more reactive than corresponding chloramines of lysine and arginine. Moreover, it can rapidly transfer chlorine to other amine groups to generate more stable chloramines (Pattison and Davies, 2005). Additionally, histidines are typically less abundant in proteins in comparison to lysines and arginines. For instance, RidA has only one histidine. Lastly, tyrosine is known to become chlorinated. However, the chlorination of tyrosine is an irreversible reaction and leads to the formation of 3-chlorotyrosine (Hawkins et al., 2003).

### LC-MS/MS analysis revealed HOCl-modified arginine, along with two modified tyrosines

Previously our laboratory reported that RidA’s full chaperone-like activity is coinciding with the addition of up to 10 chlorine atoms. However, the LC-MS/MS analysis of chlorinated RidA after tryptic digest performed in the current study only revealed one N-chlorinated amino acid, arginine R51. There was also mass spectrometric evidence for the chlorination of both tyrosines present in RidA, but, as mentioned above, this modification is irreversible and thus should not account for the reversible activation of RidA’s chaperone function. Chlorination of lysine or arginine could interfere with the tryptic digest, as these are the amino acids recognized by this particular protease. However, digest with an alternative protease, chymotrypsin did not reveal additional chlorination sites. Alternatively, N-chlorination might potentially be lost during the process of sample preparation, since the amino acid-derived monochloramines, even though less reactive, retain some of the oxidizing capacity of HOCl and are highly reactive towards oxidizable components of reaction buffers (Pattison and Davies, 2006; Ashby et al., 2020). Furthermore, N-chlorinated RidA can potentially react with sulfur-containing amino acids found in the proteases used. Therefore, a direct MS-based identification of chlorinated residues is challenging and does not result in the identification of all modified residues.

Goemans et al. conducted a similar experiment where they performed a mass spectrometric analysis of chlorinated CnoX, in which chaperone activity is also activated by HOCl. Although they identified at least 8 chlorines added to the protein mass during mass spectrometry of full-length CnoX, only five chlorinated amino acid residues were identified using LC-MS/MS analysis after protease digest, among them no lysine residue, hinting at the challenges involved in identifying this particular N-chlorinated amino acid by direct MS analysis (Goemans et al., 2018).

### Dansyl sulfinic acid can be used to stably modify N-chlorinated lysines

Searching for ways to chemically label N-chlorinated lysine, and thus making it accessible for MS-based analysis, dansyl sulfinic acid (DANSO_2_H) caught our attention. It has been previously used to derivatize low molecular weight monochloramines in water and other fluids, allowing for their detection by HPLC (Scully et al., 1984, 1986). We synthesized this probe to label the full-length RidA after its chlorination and found that it exclusively reacts with the HOCl-treated protein. We were able to detect a robust fluorescence signal once N-chlorinated RidA was treated with DANSO_2_H, but DANSO_2_H did not react with the untreated protein. Therefore, DANSO_2_H allowed for the specific labeling of N-chlorinated proteins. Using LC-MS/MS analysis of DANSO_2_H-derivatized RidA_HOCl_, we identified 6 lysine residues that we were not able to detect using a direct MS-based approach. Interestingly, these 6 lysine residues are also the lysine residues most exposed to the solvent (Table 3), suggesting they are particularly accessible to HOCl.

**Table 3.** Accessible surface area and accessibility of lysines and arginines of RidA. The analysis was performed using Accessible Surface Area and Accessibility Calculation for Protein (ver. 1.2) (http://cib.cf.ocha.ac.jp/bitool/ASA/). Relative area is the fraction of the amino acid’s total area exposed to the solvent.

| Amino acid position | Amino acid | Area (Å<sup>2</sup>) | Relative area (0.0 – 1.0) |
| --- | --- | --- | --- |
| **38** | LYS | 186.596 | 0.873 |
| **67** | LYS | 150.440 | 0.704 |
| **115** | LYS | 146.521 | 0.686 |
| **3** | LYS | 141.010 | 0.66 |
| **127** | ARG | 103.542 | 0.451 |
| **128** | ARG | 91.273 | 0.326 |
| **79** | LYS | 83.866 | 0.392 |
| **51** | ARG | 81.596 | 0.356 |
| **118** | LYS | 75.743 | 0.354 |
| **112** | ARG | 67.246 | 0.293 |
| **58** | LYS | 52.885 | 0.247 |
| **105** | ARG | 14.170 | 0.062 |
| **73** | LYS | 9.844 | 0.046 |

DANSO_2_H allows for the irreversible derivatization of monochloramines, increasing their stability and simplifying their detection using LC-MS/MS. DANSO_2_H initially reacts with monochloramines to form dansyl chloride, which then, in turn, reacts with the newly freed amino group (Scully et al., 1984; Jersey et al., 1990). The specificity of this probe in complex samples might be limited due to this issue, as there is a chance that the dansyl chloride diffuses from the originally modified amino residue. The second order rate constant of the reaction of dansyl chloride and the ε-aminogroup of lysine is 42 M^-1^ s^-^1 (Gray, 1967). Therefore, a two-step procedure with blocking of unmodified amino groups might be considered for complex samples prior to labeling with DANSO_2_H.

### Two RidA variants lacking either R105 or R128 showed a significantly decreased chaperone activity

To find out, whether specific amino acids play a key role in the activation of RidA’s chaperone-like function, all lysines and arginines were individually changed to serine residues through site-directed mutagenesis. The individual variants were then examined regarding their chaperone activity. The exchange of most lysines and arginines did not affect the chaperone activity of RidA. However, the individual exchange of amino acid residues R105 and R128 resulted in diminished chaperone activity after HOCl treatment. This could indicate that these arginines play a prominent role in chaperone activity.

Potentially, these two arginines are particularly accessible to unfolded proteins in N-chlorinated RidA and, therefore, their chlorination might be important for client protein binding. R128 is the C-terminal amino acid of RidA and, based on structural predictions, exposed to the solvent, which makes it especially accessible to unfolded proteins (Table 3). In the bacterial cytoplasm, RidA is present in the form of a trimer (Volz, 2008) and R128, R127, and K3 of every protein subunit form a large positively charged surface patch (visible in the center of the molecule in the upper panel of Fig. 7A), which is predicted to change to a more electroneutral or even negatively-charged patch after N-chlorination (Fig. 7 B, upper panel) and thus might allow for binding of unfolded client proteins.

The other residue, R105, is highly conserved in three of the seven RidA subfamilies that have enamine/imine deaminase activity (Lambrecht et al., 2012; Liu et al., 2016) and was invariable in all RidA homologs that were identified in a blast search in the SwissProt database (Altschul et al., 1990; Bateman et al., 2021) (see also Fig. 8). In *E. coli* and related species, RidA accelerates the release of ammonia from intermediates that result from the dehydration of threonine by IlvA (threonine dehydratase). Arginine 105 plays a particularly important role in the active center of the protein by forming a salt bridge with the carboxylic acid of the substrate (Lambrecht et al., 2012). This amino acid is also located in the immediate vicinity of RidA’s only cysteine at position 107. This cysteine is redox- sensitive and was modified during the nitrosative stress response (Lindemann et al., 2013). However, the presence of the cysteine is not important for RidA chaperone activity as the mutant lacking the cysteine was as active as wildtype after chlorination (Müller et al., 2014). R105, through its position in the active site, is probably also accessible to client proteins, when N-chlorinated.

We also exchanged both arginines R105 and R128 simultaneously through site-directed mutagenesis. If our argumentation that both arginines are of particular importance for client protein binding were true, we would expect the double mutant to be an even less effective chaperone than the individual mutants. Conversely, the chaperone activity of this variant corresponded approximately to the chaperone activity of the individual exchanges. One explanation would be the distant localization of these two residues, excluding a mutual influence on chaperone-like activity, even if both residues are absent. Alternatively, the lower chaperone-like activity in both single and double mutants might not be due to the lack of an N- chlorination site but the general structural disturbance that an exchange of these residues (especially in the active site of the protein) induces.

The latter possibility is further supported by the fact that the second-order rate constant of the reaction of HOCl with arginines is three orders of magnitude lower than the one for the reaction with lysines (k = 7.9×10^3^ M ^−1.^s ^−1^ vs. k = 26 M^−1.^s^−1^, respectively) (Pattison and Davies, 2001; Pattison et al., 2012). Hence, lysines are reacting faster with HOCl than arginines. Nevertheless, it is possible that these two residues are particularly reactive with HOCl. Sometimes particular amino acid residues are more reactive than suggested by the reaction rate with model compounds. As such, cysteines are usually oxidized by H_2_O_2_ with a second-order rate constant of k=14.7 ± 0.35 M^−1^s^−1^, while certain cysteines in the active site of peroxiredoxins are oxidized at a much higher velocity (k= 12’000 M^−1^s^−1^) (Peskin et al., 2013).

Therefore, it might be that R105 and R128 in RidA are particularly susceptible to N-chlorination due to the microenvironment in the protein’s structure.

### Electrostatic surface modeling of RidA_HOCl_ and the K0R0_R105S variant revealed the patterns necessary for RidA’s chaperone activity

As shown by our MS-based experiments and our mutagenesis studies, N-chlorination affects multiple basic amino acids (at least 9: lysines K3, K38, K67, K79, K115, K118 and arginines R51, R105, R128). Therefore, positive charges on the surface of a protein molecule are eliminated, causing an increase in hydrophobicity, which was previously determined experimentally using Nile Red (Müller et al., 2014; Ulfig et al., 2019) and is demonstrated by the calculated electrostatic surface potential of an N-chlorinated RidA molecule. Protein hydrophobicity as a driving force for chaperone activity is a known mechanism, and other chaperones are also known to have exposed hydrophobic surfaces that allow for the binding of client proteins (Graf et al., 2004; Mayer and Bukau, 2005; Kumar et al., 2005; Koldewey et al., 2016, 2017). A rationally designed variant, which mimics the electrostatic surface potential of N-chlorinated RidA and is indeed constitutively active as a chaperone-like holdase, substantiates this hypothesis.

## CONCLUSION

For the activation of the chaperone function of RidA, a significant increase in the protein surface hydrophobicity must take place. We present here computational and experimental evidence that this is achieved by reversible N-chlorination of nine individual lysine and arginine residues. The N-chlorination of these basic amino acids removes positive charges from the surface of RidA allowing it to bind and protect unfolded client proteins in the presence of high HOCl concentrations as they appear during inflammatory processes in bacteria-host interactions.

## MATERIALS AND METHODS

### Preparation of chlorinating agents

The concentration of NaOCl stock was determined spectrophotometrically using a JASCO V-650 UV/VIS spectrophotometer (JASCO, Tokyo, Japan) at 292 nm using the extinction coefficient ε_292_= 350 M^-1^cm^-1^. Monochloramine was prepared freshly by dropwise addition of 200 mM NaOCl solution, dissolved in 0.1 M KOH to 200 mM NH_4_Cl solution. After stirring for 5 min, the concentration of the monochloramine produced was measured spectrophotometrically using the extinction coefficient ε_242_ = 429 M^-1^cm^-1^.

### Treatment of RidA with HOCl and monochloramine (MCA)

Chlorination of RidA or RidA variants was achieved by adding 10-fold molar excess of HOCl or MCA to the protein solution. Samples were incubated for 10 min at 30 °C for HOCl-treatment and 45 min at 37 °C for MCA-treatment. Removal of the residual oxidants was achieved by size-exclusion chromatography using “Micro Bio-Spin P30 Tris Chromatography Columns” (Bio-Rad, München, Germany) or Nap-5 columns (GE Healthcare Life Sciences, Amersham, UK) according to the manufacturer’s instructions. Final protein concentrations were determined by measuring the absorbance at 280 nm using a JASCO V-650 UV/VIS spectrophotometer with an extinction coefficient of RidA of ε_280_ = 2980 M^-1^cm^-1^.

### Preparation, tryptic and chymotryptic digest of RidA_HOCl_ for mass spectrometry analysis

RidA was treated with HOCl as described above. 5 µl (approx. 25 µg of protein) of untreated and N- chlorinated RidA were mixed together with 35 µl ultrapure water and either 10 µl 5x trypsin digestion buffer (500 mM ammonium bicarbonate, pH 8.0) or 10 µl 5 x chymotrypsin digestion buffer (500 mM Tris-HCl, 10 mM CaCl_2_, pH 8.0). Freshly reconstituted trypsin in 50 mM acetic acid (1 mg/ml) or chymotrypsin in 1 mM HCl (1 mg/ml) was added to the sample for a final 1:20 enzyme-to-protein ratio.

The sample was digested at 37 °C for 18 h, and afterward, the peptides were purified using OMIX C18 tips. Purified peptides were concentrated to dryness and dissolved in 0.1 % TFA for LC-MS/MS analysis.

### Peptide cleanup using OMIX C18 tips

The samples after tryptic or chymotryptic digest were first adjusted to 0.1 % (v/v) TFA using 10 % TFA solution. Then the tips were first equilibrated 2 times with 100 µL 100 % ACN, followed by 2 times washing with 100 µL 50 % ACN, 0.1 % TFA, and lastly 2 times washing with 0.1 % TFA. Afterwards, the sample was loaded onto the tip by pipetting it 8 times slowly up and down without releasing the pipette. The tip was then washed 3 times in 500 µL 0.1 % TFA and the sample was eluted in 40 µL 75 % ACN, 0.1 % TFA by pipetting it 8 times.

### Liquid chromatography-mass spectrometry (LC-MS/MS)

Dansyl-derivatized or unmodified peptides after tryptic or chymotryptic digest were analyzed via LC- MS/MS with an LTQ Orbitrap Elite (Thermo Fisher Scientific) as follows: 100 ng of the sample were loaded onto a C18 precolumn (100-µm × 2-mm Acclaim PepMap100, 5 µM, Thermo Fisher Scientific) with 2.5 % ACN/0.1 % TFA (v/v) at a flow rate of 30 µl/min for 7 min. The peptides were then loaded onto the main column (75-µm × 50-cm Acclaim PepMap100 C18, 3 µm, 100-Å, Thermo Fisher Scientific) with 95 % solvent A (0.1 % formic acid (v/v)) and 5 % solvent B (0.1 % formic acid, 84 % ACN (v/v)) at a flow rate of 0.4 µl/min. Peptides were eluted with a linear gradient of 5–40 % B (120 min, 0.4 µl/min). The 20 most intense peaks were selected for MS/MS fragmentation using collision-induced dissociation in the linear ion trap (charge range +2 to +4, exclusion list size: 500, exclusion duration: 35 s, collision energy: 35 eV).

The generated raw files after LC-MS/MS were analyzed using Xcalibur Software (Qual Browser 3.1, Thermo Fischer Scientific, Waltham, MA, USA).

### Mass spectrometry data analysis using MaxQuant

MaxQuant software (version 1.5.1.0, DE) was used to identify and quantify dansyl- and chlorine- modified peptides. The following modifications were added to the variable modification list of the search engine Andromeda: dansylation (+233.05, lysine, arginine, and histidine), chlorination (+33.96, lysine, arginine, histidine, tyrosine) and trioxidation (+47.98, cysteine). For peptide search using Andromeda, the *E. coli* K12 proteome database (taxonomy ID83333) obtained from UniProt (4518 proteins, released September 2019, The UniProt Consortium, 2019) was used. Two miscleavages were allowed, Oxidation (M), Dansylation (KR), Chlorination (KRYH), Carbamidomethyl (C), Trioxidation (C) were chosen as variable modification. Identified peptides were assessed using the “peptides.txt” MaxQuant output file. The availability of modification was monitored using “dansylation-sites.txt”, “chlorination-sites.txt” and “modificationSpecificPeptides.txt” output files. Data was imported and analyzed using Microsoft Excel Version 16.20 (Microsoft, Redmond, WA, USA). All peptide modifications were verified by manual examination of MS/MS spectra using the criteria proposed by Nybo et al., 2018. Described criteria are: 1) identification of unmodified peptide; 2) coverage of modification site by fragment ion series; 3) correct assignment of peaks in MS/MS spectra (neutral losses: Met + O (−64), Met + 2O (−80), Cys + O (−50), Cys + 2O (−66), Cys + 3O (−82)); 4) similar fragmentation patterns between modified and unmodified peptide(s).

### Construction of RidA variants with single amino acid substitutions

*E. coli* strains, plasmids, and primers used in this study are listed in table 4. Single nucleotides in the *ridA* gene were exchanged using PCR-based mutagenesis, known as QuickChange PCR. PCR was performed using 150 ng of pUC19_*ridA* or pEX_ridA-K0R0 as a template and 125 ng of each specific primer (Table 5) for every single exchange. 20 µL of the PCR product were digested with 20 U of DpnI at 37 °C for 1 h to eliminate the template plasmid. Subsequently, *E. coli* XL-1 blue cells were transformed with the sample using a standard heat-shock method and plated on LB agar plates supplemented with 100 mg/ml ampicillin. Plasmid DNA was isolated from single colonies, and successful mutagenesis was verified by sequencing. Afterwards, *ridA* gene variants were subcloned into pET22b(+) expression vector via the restriction sites *Nde*I and *Xho*I. The resulting pET22b(+)-based constructs were transformed into *E. coli* BL21(DE3) cells for the subsequent overexpression of RidA variants.

**Table 4:**
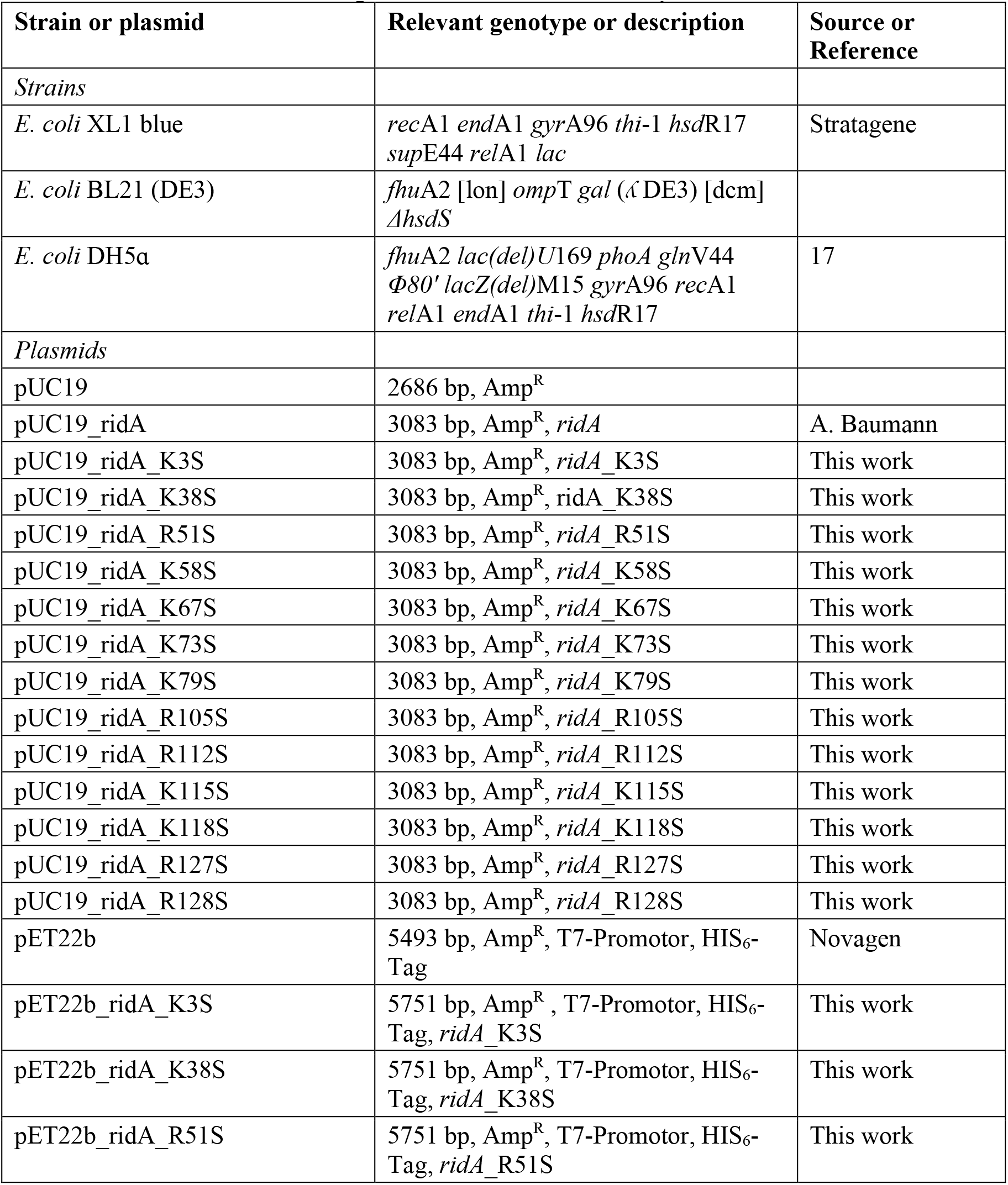
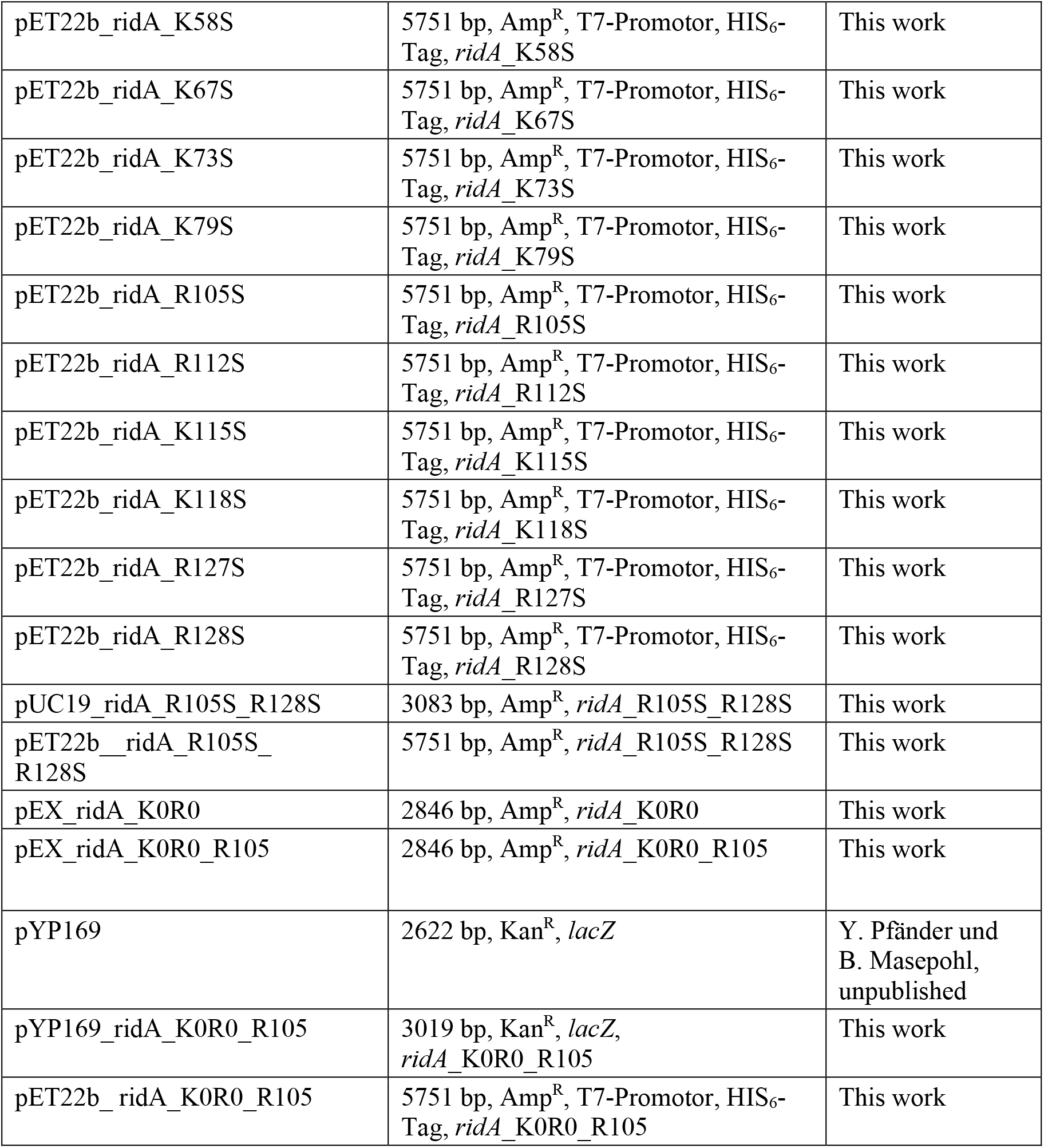
Bacterial strains and plasmids used in this study.

**Table 5:**
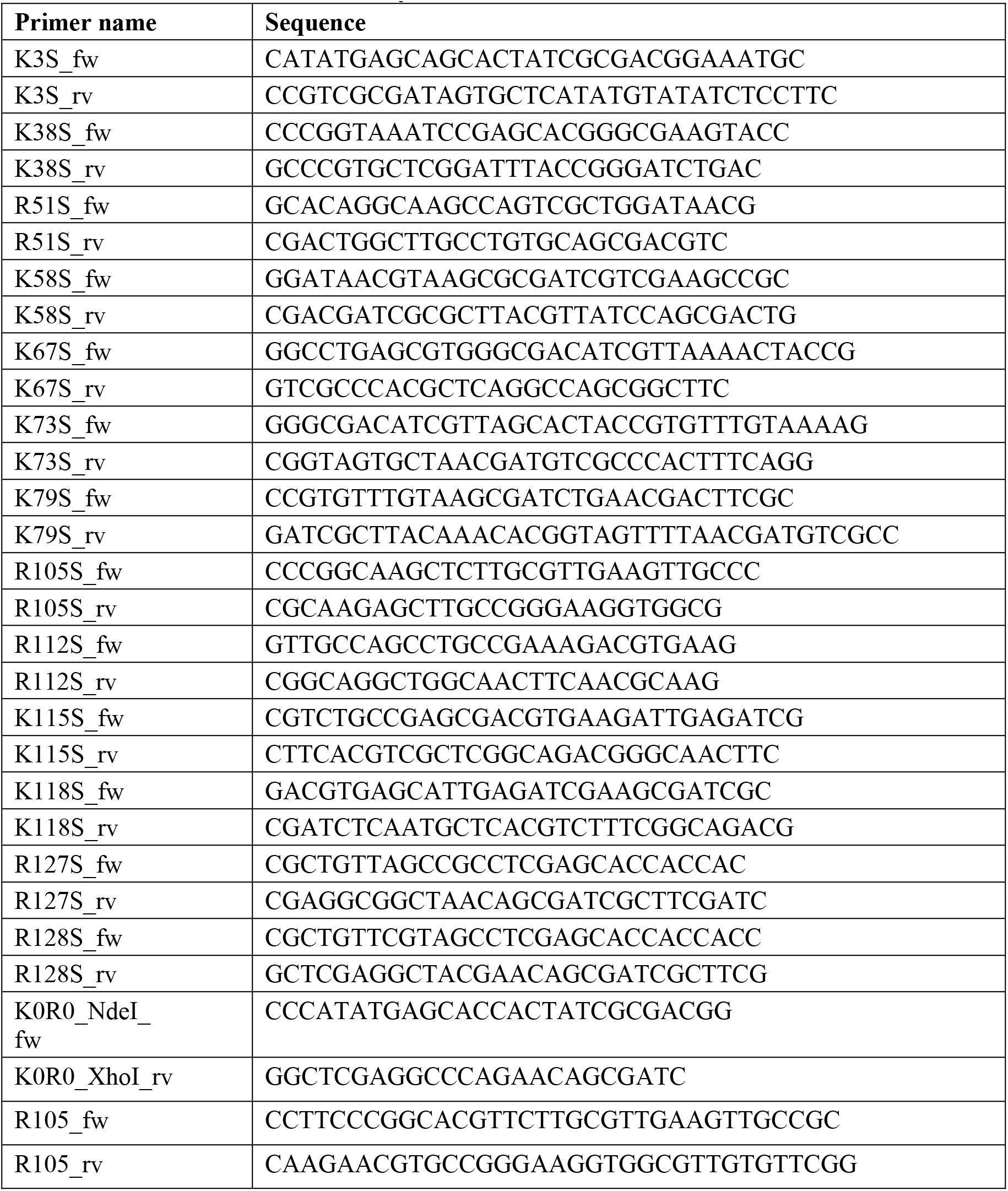
Primers used in this study.

### Overexpression and purification of RidA variants

For overexpression, a single colony of *E. coli* BL21 (DE3) carrying the respective pET22b(+) plasmid was inoculated in 50 mL LB containing 200 mg/L ampicillin and grown overnight at 37 °C, 120 rpm. The next morning, the overnight culture was used to inoculate 5 L of LB medium, supplemented with ampicillin, and incubated at 37 °C and 120 rpm until the OD_600_ reached 0.5-0.6. At this point, protein expression was induced by the addition of 1 mM isopropyl 1-thio-ß-D-galactopyranoside (IPTG) to the culture. After 3 h of incubation at 37 °C, 120 rpm, cells were harvested by centrifugation at 7800 x g and 4 °C for 45 min and either stored at -80 °C or directly used for the purification procedure.

The resulting cell pellet was washed once with lysis buffer (50 mM sodium phosphate, 300 mM NaCl, 10 mM imidazole, pH 8.0) containing 1 ml of EDTA-free protease inhibitor mixture (Roche Applied Science). Cells were disrupted by passing the cell suspension three times through a Constant systems cell disruption system TS benchtop device (Score Group plc, Aberdeenshire, UK) at 1.9 kbar and 4 °C, followed by the addition of PMSF to a final concentration of 1 mM.

The cell lysate was centrifuged at 6700 x g, 4 °C for 1 h, and the supernatant was vacuum filtered through a 0.45 μm filter. The filtrate was loaded onto a Ni-NTA affinity column. The column was washed with 10 mL washing buffer (50 mM sodium phosphate, 300 mM NaCl, 20 mM imidazole, pH 8.0). For purification of the R0K0_R105 variant, washing buffer was additionally supplemented with 1 M NaCl and 0.1 % SDS. Purified proteins were stored at -80 °C and K0R0_R105 was stored at room temperature. Protein concentrations were determined using a JASCO V-650 spectrophotometer using the extinction coefficient ε_280_=2,980 M^-1^cm^-1^.

### Protein aggregation assay with citrate synthase

Citrate synthase was chemically denatured in 4.5 M GdnHCl, 40 mM HEPES, pH 7.5 at room temperature overnight. The final concentration of denatured citrate synthase was 12 mM.

To monitor initial aggregation of citrate synthase, 20 µl of denatured protein were added to 1580 µl of 40 mM HEPES, pH 7.5 to a final concentration of 0.15 µM after 20 s of measurement. To test the chaperone activity, RidA or RidA variants were added prior to the addition of citrate synthase to the buffer at different molar access (0.5-16-fold) over dimeric citrate synthase. The increase in light scattering was monitored for 240 s using a JASCO FP-8500 fluorescence spectrometer equipped with an EHC-813 temperature-controlled sample holder (JASCO, Tokyo, Japan). Measurement parameters were set to 360 nm (Em/Ex), 30 °C, medium sensitivity, slid width 2.5 nm (Em/Ex). Relative chaperone activity of different RidA variants was calculated based on the difference between initial and final light scattering. The chaperone activity of wild-type chlorinated RidA was set to 100 %.

### Synthesis of N,N-Dimethyl-1-amino-5-naphtalenesulfinic acid (Dansyl sulfinic acid, DANSO_2_H)

Dansyl sulfinic acid (DANSO_2_H) was synthesized from dansyl chloride, as described in Scully et al., 1984 with minor modifications. Dansyl chloride (5 g) was added to a continuously stirred aqueous solution of sodium sulfite (10.7 g in 50 mL) warmed to 70 °C. The reaction temperature was kept at 80 °C for 5 h. After the solution was cooled, DANSO_2_H was precipitated from the product mixture by acidifying the solution to pH 4 with concentrated sulfuric acid. The precipitate was then filtered. The precipitate was then dried in a vacuum desiccator over silica gel. The powder was re-dissolved in a cold, aqueous solution of NaOH (2.8 M). The resulting solution was filtered and titrated to pH 4.0 using sulfuric acid. The resulting DANSO_2_H was dried again in a vacuum desiccator and stored in the light- protected vial at 4 °C. The purity of the resulting dansyl sulfinic acid (retention time 6.48 min) was confirmed by HPLC (Fig. S1) and was 95 %. The 5 % contamination by a corresponding sulfonic acid (retention time 6.55 min) does not affect the derivatization of chloramines (Scully et al., 1984; Alpmann and Morlock, 2008).

### Derivatization of RidA with dansyl sulfinic acid and dansyl chloride

200 mM DANSO_2_H solution was prepared freshly by dissolving the powder in 200 mM NaHCO3 buffer, pH 9.0, and 10 % (w/w) DANSCl stock solution (370 mM) (abcr, Karlsruhe, Germany) in acetone was directly used.

RidA was chlorinated with HOCl as described above. Using an NAP-5 gel filtration column, residual HOCl was removed, and the buffer was exchanged to 200 mM NaHCO_3_ buffer, pH 9.0. 250 µM RidA_HOCl_ or RidA_UT_ were incubated with a 50-fold molar excess of DANSCl or DANSO_2_H for 1h, 37 °C, 1300 rpm. The derivatizing agent was removed using an NAP-5 gel filtration column. Successful derivatization of RidA by monitoring the fluorescence of the resulting sulfonamide was determined by a fluorescence emission scan from 360–600 nm in a JASCO FP-8500 fluorescence spectrometer with the following parameters: 340 nm excitation, 2.5 nm slit width (Ex/Em) and medium sensitivity.

### Preparation and chymotryptic digest of dansylated proteins for mass spectrometry analysis

5 µL (approx. 8 µg of protein) of each dansylated sample prepared above were mixed together with 25 µl ultrapure water and 10 µl 5 x chymotrypsin digestion buffer (500 mM Tris-HCl, 10 mM CaCl_2_, pH 8.0). pH of the resulting solution was monitored to be around 8.0. Then, DTT was added to the solution to a final concentration of 10 mM and incubated at 60 °C for 45 min. After the sample was cooled to room temperature, 3.5 µL of freshly prepared 500 mM iodoacetamide solution in pure water were added to a final concentration of 20 mM. The sample was incubated at room temperature for 30 min, protected from light. To quench the alkylation reaction, 1 ul of 500 mM DTT was added, and the volume of the solution was adjusted to 50 µl using pure water. Chymotrypsin, reconstituted to a concentration of 1 mg/mL in 1 mM HCl, was then added to the sample for a final 1:20 enzyme-to-protein ratio. The reaction mixture was incubated at 37 °C for 18 h. The sample was then desalted using OMIX C18 tips (Agilent Technologies, USA) according to the manufacturer’s instructions. Purified peptides were concentrated to dryness and dissolved in 0.1 % TFA for LC-MS/MS analysis.

### Protein structure modification and electrostatic surface modelling of RidA, RidA_HOCl_, and K0R0_R105

*E. coli* RidA crystal structure was accessed using PDB entry number 1QU9 and visualized using PyMOL 2.3.4 (Schrödinger, New York, NY, USA). This structure was converted to a PQR file using the PDB2PQR webserver (Jurrus et al., 2018) under an assumed pH of 7.0, using PROPKA to assign protonation and the AMBER forcefield to assign partial charges and atomic volumes to the structure’s atoms. Using the resulting PQR file as input, PyMol’s APBS plugin was used to calculate the electrostatic surface potential.

To model the N-chlorinated RidA_HOCl_, the PQR file of RidA was manipulated using a custom python script (Supplemental Material S2). This script removes charge bearing protons from specified lysine and arginine residues, rectifies binding angles (in case of deprotonated arginine) and replaces one nitrogen- attached hydrogen atom in these residues with a chlorine atom using the following bond geometries and partial charges: Charge of the chlorine atom: 0.07, charge of the nitrogen atom: -0.61 (based on values for CH_3_NHCl (Heeb et al., 2017)), radius of the chlorine atom 1.75 Å (Bondi, 1964), length of the N-Cl bond: 1.784 Å (based on values for NH_2_Cl (Harmony et al., 1979)). The resulting manipulated PQR file was then used to calculate the electrostatic surface potential of RidA_HOCl_ using the APBS plugin of PyMol.

The structure of RidA variant R0K0_R105 was modeled using the mutagenesis function of PyMOL. The specified amino acids in wild-type RidA were exchanged as follows:

K3 → T3

K38 → A38

R51 → H51

K58 → G58

K67 → S67

K73 → S73

K79 → T79

R112 → A112

K115 → L115

K118 → L118

R127 → L127

R128 → G128

The electrostatic surface potential for this variant was then calculated as outlined for the wildtype above (conversion of the pdb-file to a PQR file using the PDB2PQR webserver under an assumed pH of 7.0, using PROPKA to assign protonation and the AMBER forcefield and subsequent use of the ABPS plugin in PyMol).

## ACKNOWLEDGMENTS

LIL acknowledges funding from the DFG Priority Program 1710 “Dynamics of Thiol-based Redox Switches in Cellular Physiology” through grant LE2905/1-2. LIL, EH, and JEB would like to thank the DFG Research Training Grant 2341 “Microbial Substrate Conversion” for supporting this work, and JEB further acknowledges funding from the DFG CRC1316-1 and BA 4193/7-1.

## AUTHOR CONTRIBUTIONS

MV, JF, AM, LIL conceived and designed the study. NL assisted in and performed protein expression and purification experiments. MV performed mass spectrometry and fluorescence spectroscopy experiments and evaluated MS data. MV and LIL synthesized and characterized the DANSO_2_H probe. YS and KSC designed and assisted in the synthesis of the DANSO_2_H probe. JF designed and performed site directed mutagenesis experiments, mass spectrometry experiments, and citrate synthase aggregation assays. AM designed site directed mutagenesis experiments and the K0R0_R105 variant and performed citrate synthase aggregation assays. KR and BS performed mass spectrometry measurements and assisted in MS data handling. MK, CJ, TJ, EH, JEB consulted on the computational structure prediction and calculation of the electrostatic surface potential of RidA, RidA_HOCl_ and K0R0_R105. LIL and MV calculated the electrostatic surface potentials. MV, AM, JF, LIL wrote the manuscript, all other authors consulted on the manuscript.

## SUPPLEMENTAL MATERIAL

**Figure S1.**
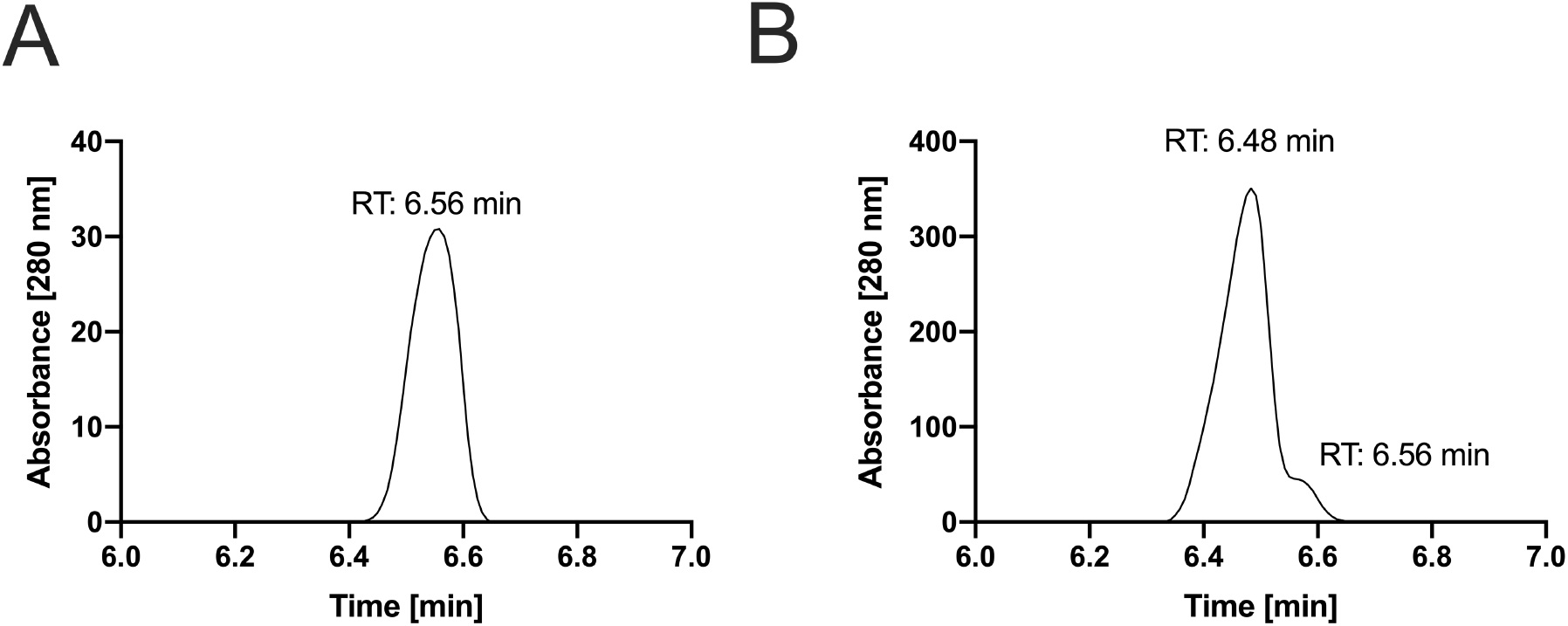
High-performance liquid chromatography (HPLC) of DANSO3H and DANSO2H. (**A)** HPLC measurement of 10 µM dansyl sulfonic acid (DANSO3H). The absorbance A280 reached its maximum at 6.56 min. (**B)** HPLC of 100 µM dansyl sulfinic acid (DANSO2H). Two absorbance peaks could be distinguished: at 6.48 min, corresponding presumably to DANSO2H and a lower peak at 6.56 min corresponding to the contamination with DANSO3H, as determined using DANSO3H-standard.

**Suppelemental Material S2:**
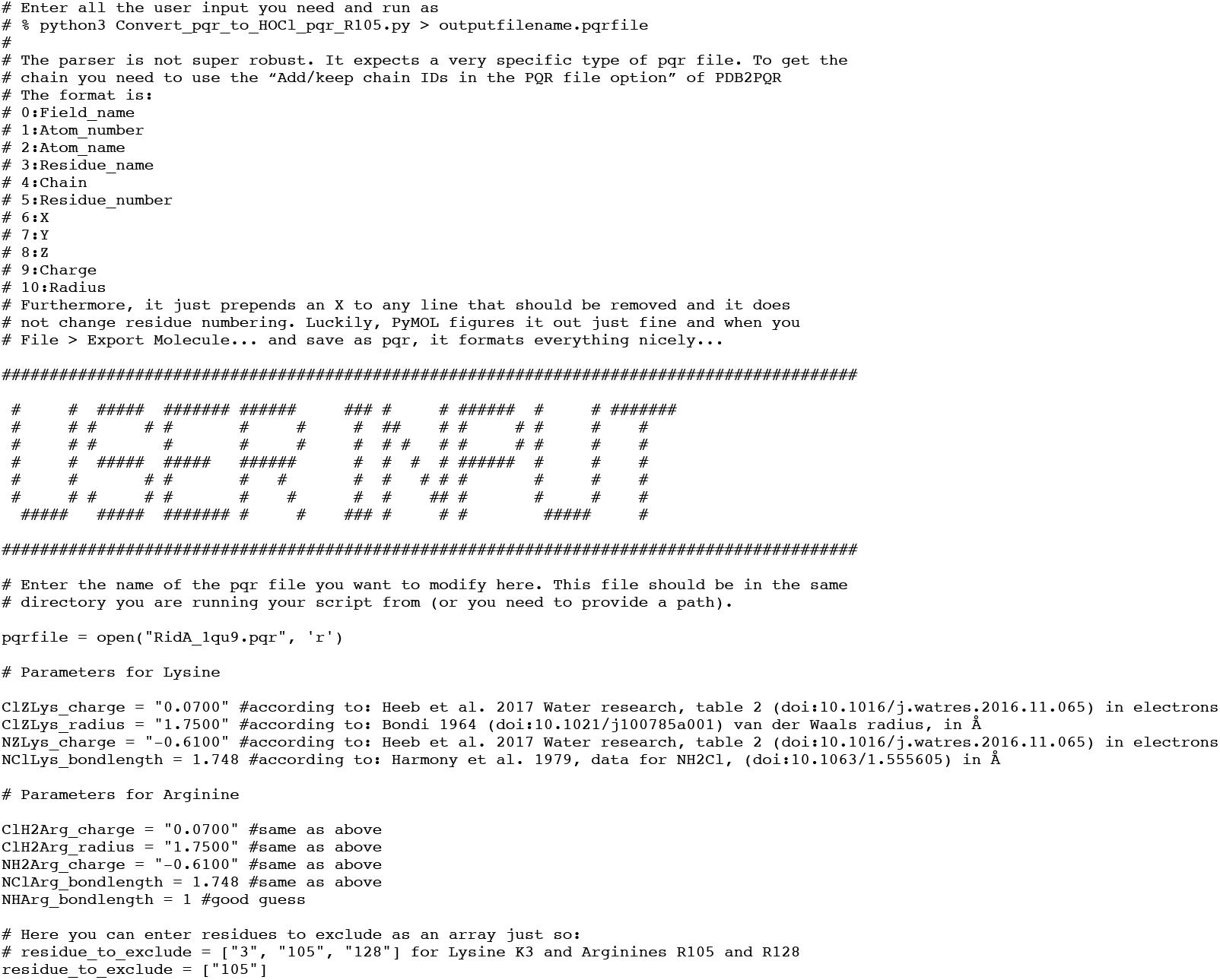
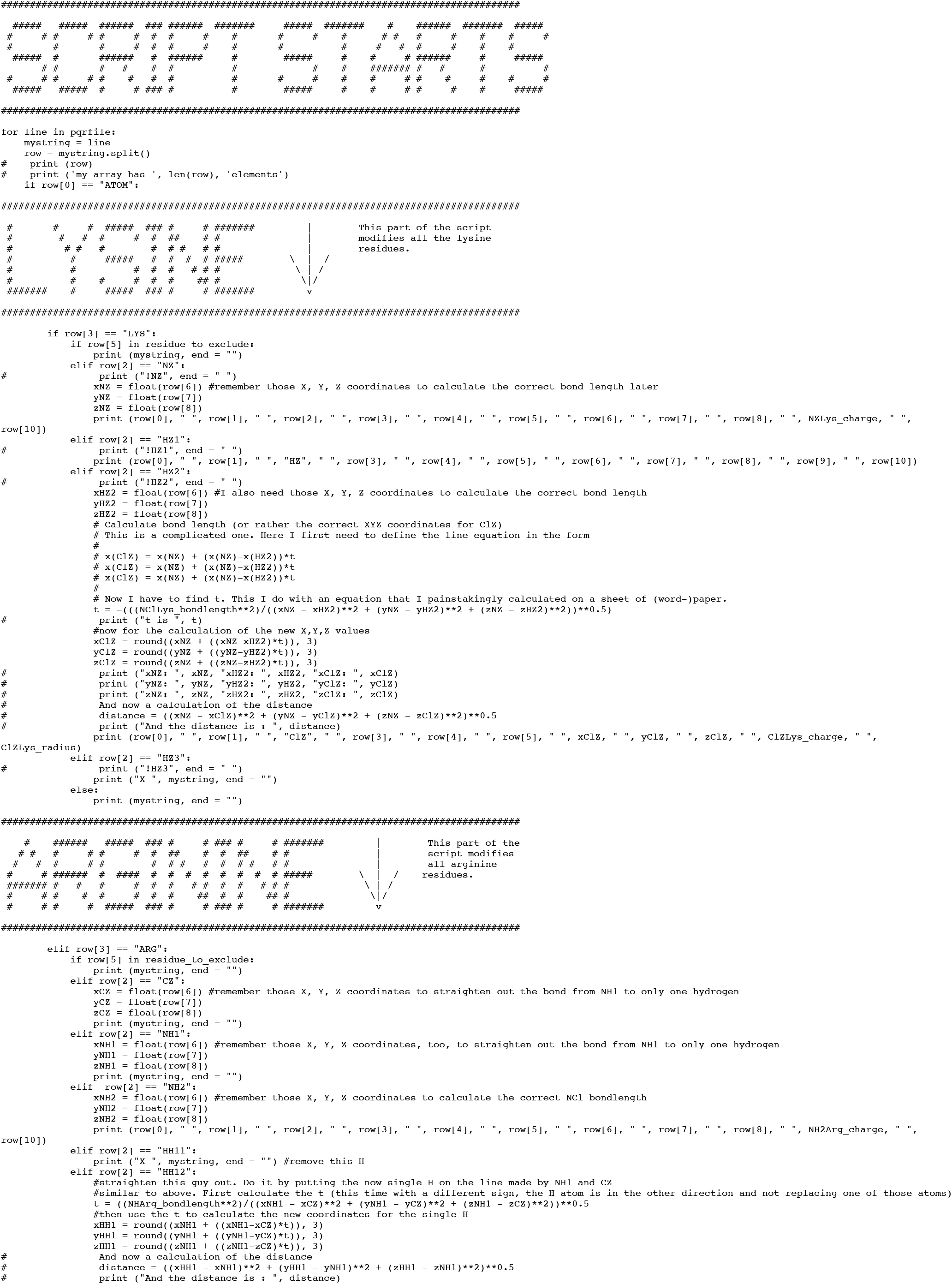
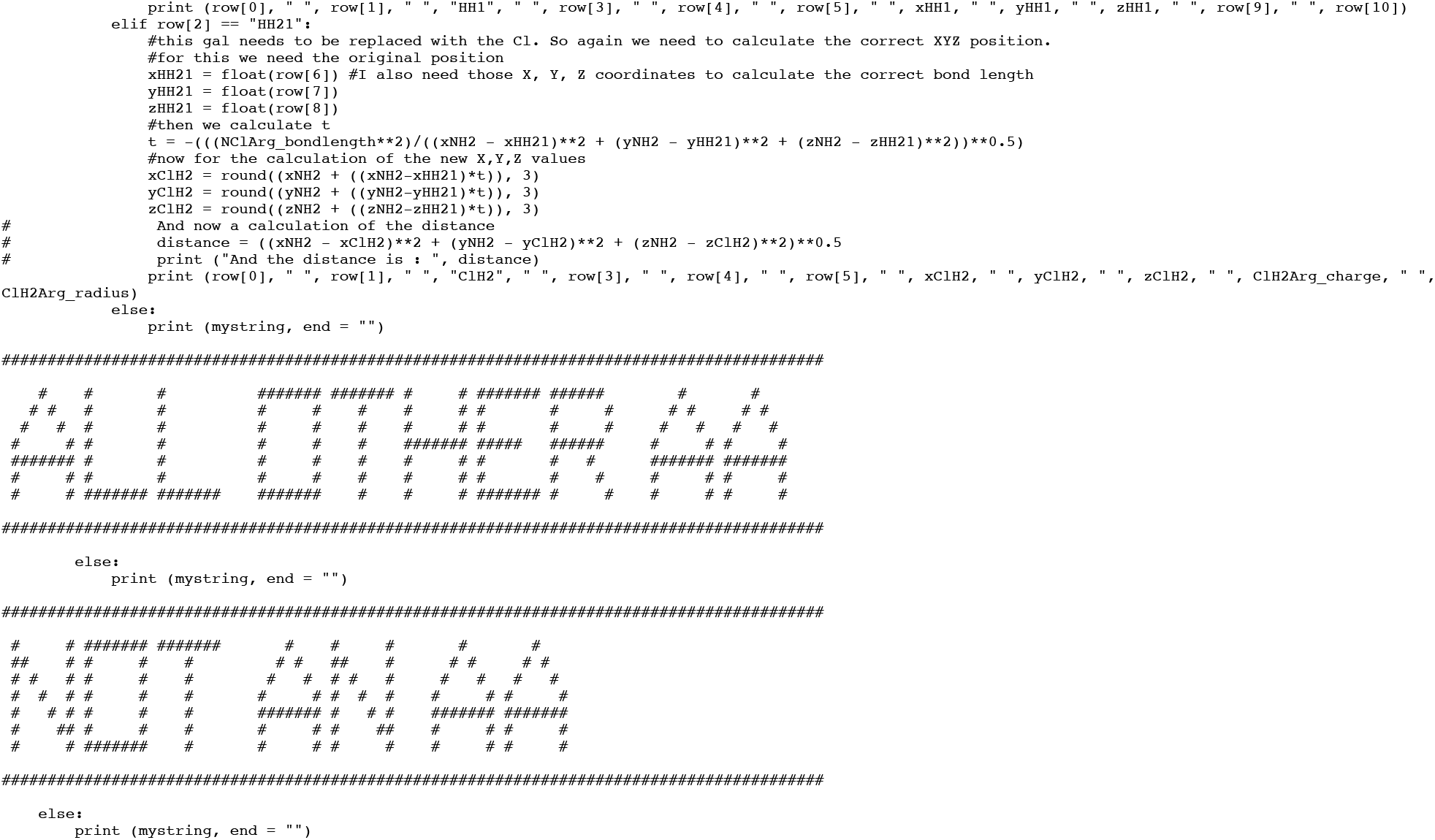
**Python script for the manipulation of PQR-files to generate a predicted structure for RidAHOCl from PDB structure 1qu9.**

